# CDK1 dependent phosphorylation of hTERT contributes to cancer progression

**DOI:** 10.1101/556514

**Authors:** Mami Yasukawa, Yoshinari Ando, Taro Yamashita, Yoko Matsuda, Shisako Shoji, Masaki S. Morioka, Hideya Kawaji, Kumiko Shiozawa, Takaya Abe, Shinji Yamada, Mika K. Kaneko, Yukinari Kato, Yasuhide Furuta, Tadashi Kondo, Mikako Shirouzu, Yoshihide Hayashizaki, Shuichi Kaneko, Kenkichi Masutomi

**Author notes:** These authors contributed equally to this work (Mami Yasukawa, Yoshinari Ando, Taro Yamashita).

## Abstract

The telomerase reverse transcriptase is upregulated in the majority of human cancers and contributes directly to cell transformation. Here we report that hTERT is phosphorylated at threonine 249 during mitosis by the serine/threonine kinase CDK1. Clinicopathological analyses revealed that phosphorylation of hTERT at threonine 249 occurs more frequently in advanced cancers. Using CRISPR/Cas9 genome editing, we introduced substitution mutations at threonine 249 in the endogenous *hTERT* locus and found that phosphorylation of threonine 249 is necessary for hTERT-mediated RNA dependent RNA polymerase (RdRP) activity but dispensable for reverse transcriptase activity. Cap Analysis of Gene Expression (CAGE) demonstrated that hTERT phosphorylation at 249 regulates the expression of specific genes that are necessary for cancer cell proliferation and tumor formation. These observations indicate that phosphorylation at threonine 249 regulates hTERT RdRP and contributes to cancer progression in a telomerase independent manner.

## Introduction

Recent analyses of human cancer genomes revealed that somatic mutations within the promoter of human telomerase reverse transcriptase (*hTERT*) occur at high frequency in over 50 distinct cancer types (*1-6*). These common promoter mutations are associated with higher expression levels of *hTERT* and a poor prognosis (*5,7-9*). In humans, experiments involving live cell imaging techniques combined with CRISPR-Cas9 genome editing demonstrated that recruitment of hTERT and *hTERC* to telomeres occurs through dynamic interactions between telomerases and the chromosome end during S-phase (*10*). Although these observations indicate that recruitment of telomerase holoenzyme to the telomere is regulated in cell-cycle dependent manner, only a small subset of hTERT forms interactions with telomeres and Cajal bodies even in S-phase (*10*) and the regulation and function of the majority of hTERT outside S-phase is poorly understood.

## Results

### Mitotic specific accumulation of hTERT

Since it has been challenging to detect endogenous hTERT (*11,12*), we extensively validated available hTERT-specific antibodies against hTERT including the mouse monoclonal antibody (mAb) (clone 10E9-2), the mouse mAb (clone 2E4-2), the sheep polyclonal Abs (pAbs) abx120550 and the rabbit mAb ab3202. Specifically, we performed validation experiments by (i) immunoprecipitation (IP) with anti-hTERT antibodies followed by immunoblotting (IB) (**Fig. 1A**), (ii) suppression of hTERT by siRNAs specific for *hTERT* (*13,14*) followed by IP-IB (**Fig. S1A**) and (iii) recovery of telomerase activity with these antibodies as gauged by direct telomerase assay (*12,15*) (**Fig. S1B**). In each case, we found that the 10E9-2, the 2E4-2, the abx120550 and the ab32020 identified hTERT (**Fig. 1A** and **Fig. S1**), confirming the sensitivity and specificity of these antibodies.

**Fig. 1.**
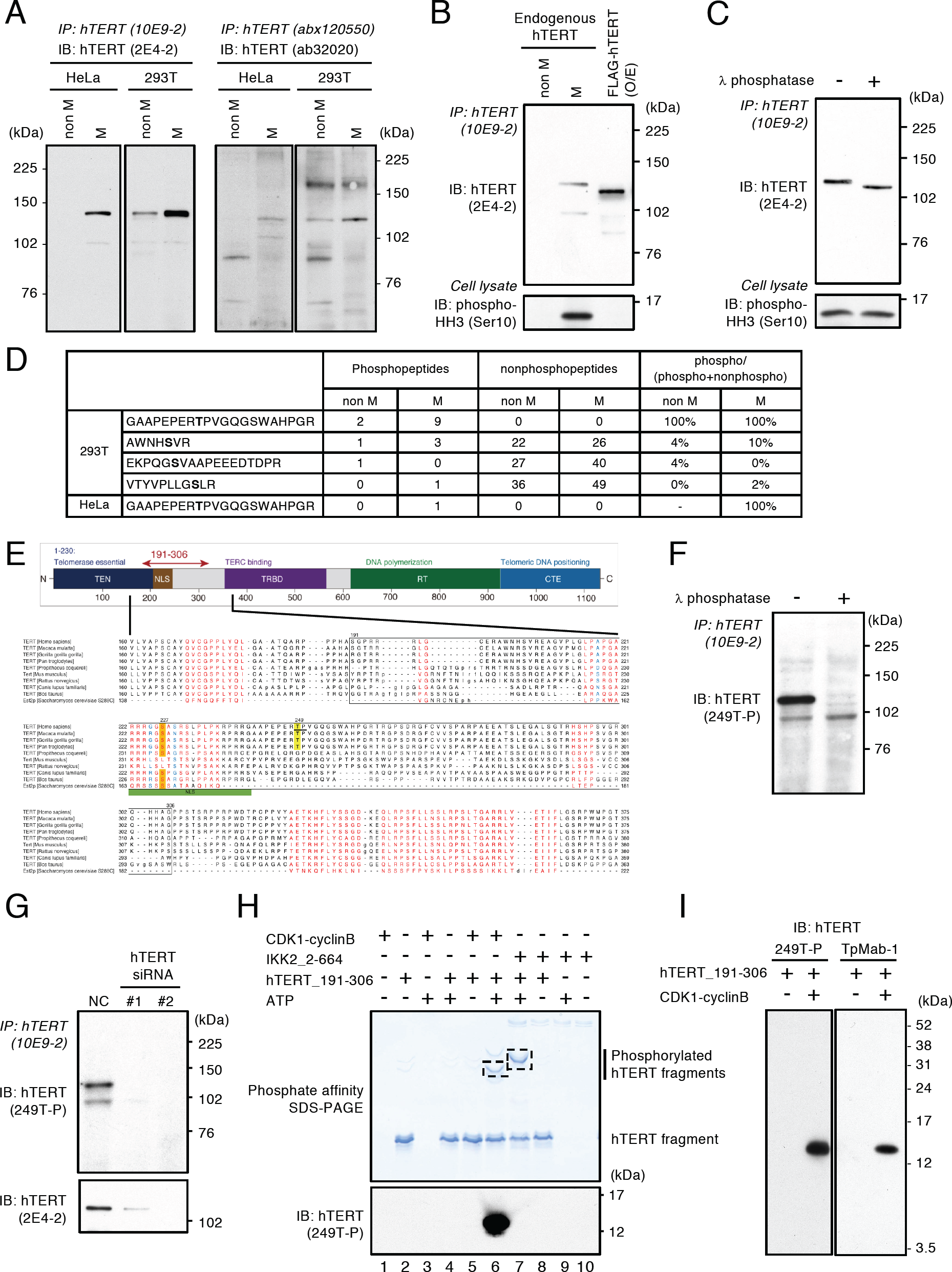
Identification of phosphorylation site of hTERT in mitosis. **(A)** Detection of endogenous hTERT in HeLa and 293T cells. The cells were treated with DMSO (denoted as “non M”) or nocodazole to manipulate cells in mitotic phase (denoted as “M”), and immunoprecipitated with anti-hTERT mouse mAb (clone 10E9-2) or anti-hTERT sheep pAbs (abx120550). The hTERT proteins were detected by anti-hTERT mouse mAb (clone 2E4-2) (two left panels) or anti-hTERT rabbit mAb (ab32020) (two right panels). **(B)** IP-IB of endogenous hTERT and FLAG-tagged hTERT. HeLa cells were treated with DMSO (non M) or nocodazole (M). HeLa cells expressing FLAG-tagged hTERT were treated with DMSO. hTERT proteins were isolated by immunoprecipitation with anti-hTERT mAb (clone 10E9-2) and detected by anti-hTERT mAb (clone 2E4-2) (upper panel). Cells arrested in mitosis with nocodazole were confirmed by anti-phospho-histone H3 (Ser10) antibodies (lower panel). **(C)** IP-IB of endogenous hTERT using 293T cells synchronized to mitotic phase. The immune complexes were treated with *λ* phosphatase to remove the phosphate groups from hTERT proteins and detected by anti-hTERT mAb (clone 2E4-2) (upper panel). Accumulation in mitotic phase was confirmed by anti-phospho-histone H3 (Ser10) antibodies (lower panel). **(D)** Summary of MS analysis to identify phosphorylation sites in hTERT. The numbers of phosphopeptides and nonphosphopeptides are based on pep_expect (Expectation value for PSM) <0.05 used as a cutoff value. Bold letters indicate amino acid residues identified as phosphorylated residues. See the details in **Supplementary Data S1**. **(E)** Multiple sequence alignment of telomerases using COBALT (*63*). Red letters denote conserved residues and blue letters denote non-conserved residues with no gaps. Threonine 249 highlighted with yellow is the phosphorylated threonine residue detected in this study. Serine 227 highlighted with orange is known to be phosphorylated by Akt in previous study (*17*). **(F)** Detection of endogenous hTERT immunoprecipitated from HeLa cells manipulated in mitosis. The immune complexes were treated with *λ* phosphatase. Phosphorylation of hTERT at T249 was detected by anti-249T-P rabbit pAbs. **(G)** IP-IB of endogenous hTERT proteins using HeLa cells transfected with two different siRNAs specific for *hTERT* followed by nocodazole treatment. Phosphorylation of hTERT at T249 was detected by anti-249T-P pAbs (upper panel). Whole hTERT proteins were detected by anti-hTERT mAb (clone 2E4-2) (lower panel). **(H)** Phosphate affinity SDS-PAGE separation of phosphorylated hTERT_191-306 proteins. The recombinant hTERT fragment proteins were phosphorylated by CDK1-cyclinB or IKK2_2-664 *in vitro* and phosphorylated proteins were detected in Phos-Tag SDS-PAGE (upper panel, lanes 6 and 7). Phosphorylation of hTERT at threonine 249 by CDK1-cyclinB was confirmed by anti-249T-P pAbs (lower panel). The protein bands enclosed with square were analyzed by MS to confirm the phosphorylation sites. **(I)** Phosphorylation of threonine 249 by CDK1-cyclinB was confirmed by anti-249T-P pAbs and TpMab-1 mouse mAb.

We previously reported that hTERT protein localizes to mitotic spindles (*16*) and hTERT protein expression is enriched in mitosis (*14*) (**Fig. 1A**). We reconfirmed the enrichment of hTERT protein in mitosis using the antibodies described above (*12*) (**Fig. 1A**, two right panels). In addition, we confirmed mitotic specific accumulation of hTERT at the mRNA level (**Fig. S2**). These observations suggest that expression of hTERT protein is regulated in a cell-cycle dependent manner.

### Phosphorylation of hTERT in mitosis

To investigate hTERT regulation in mitosis, we first treated HeLa cells with nocodazole. We confirmed that cells accumulated in mitotic phase by assessing phospho-histone H3 (Ser10) levels (**Fig. 1B**, lower panel). When we examined the migration of endogenous hTERT in the mitotic phase by SDS-PAGE, we found that endogenous hTERT isolated by immunoprecipitation with anti-hTERT mAb (clone 10E9-2) (*16*) migrated slower than ectopically expressed FLAG-tagged hTERT (**Fig. 1B**, upper panel). We thus speculated that endogenous hTERT in mitotic phase is post-translationally modified. We treated hTERT immunoprecipitated with anti-hTERT mAb (clone 10E9-2) from mitotic cells with *λ* phosphatase and found that phosphatase treatment diminished the mobility shift of hTERT protein (**Fig. 1C**). This observation suggested that hTERT is phosphorylated in mitosis.

To identify the mitotic phosphorylation sites in hTERT, we ectopically expressed hTERT in HEK-293T (293T) or HeLa cells followed by treatment with nocodazole to arrest cells in mitosis. We isolated hTERT by immunoprecipitation and performed mass spectrometry (MS) analysis using liquid chromatography-tandem mass spectrometry (LC-MS/MS). We used the Mascot software package (version 2.5.1; Matrix Science) to search for the mass of each peptide ion peak against the SWISS-PROT database (Homo sapiens, 20,205 sequences in the Swiss prot_2015_09.fasta file) and to identify the phosphorylation sites. We identified four phosphopeptides in 293T cells and one phosphopeptide in HeLa cells. The phosphopeptide ^241^GAAPEPERpTPVGQGSWAHPGR^261^ was found in both 293T and HeLa cells (**Fig. 1D**). Moreover, phosphorylation sites analysis by the Mascot software revealed that only the residue at threonine 249 (T249) was phosphorylated in the phosphopeptide (amino acids 241-261) (**Supplementary Data S1**). We found that T249 is conserved in primates, while serine 227, corresponding to nuclear localization of hTERT by the Akt-mediated phosphorylation (*17*), is conserved among most mammals (**Fig. 1E**).

To interrogate the consequences of hTERT phosphorylation at T249, we generated rabbit polyclonal antibodies that specifically recognize phosphothreonine 249 (anti-249T-P). We confirmed that treatment of hTERT isolated from mitotic HeLa cells with *λ* phosphatase or suppression of hTERT by hTERT-specific siRNAs ablated the signal detected by this antibody (**Figs. 1F** and **1G**). Taken together, these observations indicate that hTERT at T249 is phosphorylated in mitosis.

We noted that the sequence surrounding T249 is part of a motif that is frequently phosphorylated in mitosis and has been implicated as a target of the serine-threonine kinase cyclin-dependent kinase 1 (CDK1) (*18*). To investigate whether CDK1 phosphorylates hTERT, we performed *in vitro* kinase assay using recombinant hTERT fragment proteins (amino acids 191-306) and CDK1-cyclinB proteins followed by MS analysis. We used phosphate-affinity gel electrophoresis (Phos-tag SDS-PAGE) to detect the phosphorylation status of hTERT fragment incubated with purified CDK1-cyclinB or IKK2_2-664 used as a control of serine-threonine kinase. Phos-Tag SDS-PAGE allows the migration delay of phosphorylated proteins relative to their unphosphorylated counterparts in gels containing a phosphate-binding tag (*19*). The Phos-tag SDS-PAGE revealed that hTERT_191-306 proteins were phosphorylated by CDK1-cyclinB and IKK2_2-664 *in vitro* (**Fig. 1H**, upper panel). The hTERT_191-306 protein phosphorylated by CDK1-cyclinB was detected with the anti-249T-P antibodies, but we were unable to detect the protein phosphorylated by IKK2_2-664 with the anti-249T-P antibodies (**Fig. 1H**, lanes 6 and 7 on lower panel), suggesting that IKK2 does not phosphorylate hTERT at T249 *in vitro*. To further confirm that hTERT T249 is phosphorylated by CDK1-cyclinB *in vitro*, we performed MS analysis using LC-MS/MS and identified that threonine 249 was phosphorylated by CDK1-cyclinB, not by IKK2_2-664 (**Fig. S3** and **Table S1**). These observations confirmed that CDK1-cyclinB phosphorylates hTERT T249.

To further study hTERT phosphorylation at T249, we generated a mouse monoclonal antibody that specifically recognize phosphothreonine 249 (TpMab-1). We confirmed that the antibody specifically detected phosphorylation signals of hTERT T249 phosphorylated by CDK1-cyclinB *in vitro* similar to anti-249T-P polyclonal antibodies (**Fig. 1I**).

### Phosphorylation of hTERT at T249 occurs in cancers from clinical samples

To assess whether T249 was phosphorylated in human cancers, we first validated the specificity of the anti-249T-P pAbs for immunohistochemical (IHC) staining in human tissue sections. The observed signals were fully abolished by the absorption treatment with phosphopeptide (**Fig. S4**), whereas treatment with nonphosphopeptide did not alter the signals (**Fig. S4**), indicating that the antibodies are specific and applicable for immunohistochemical analyses.

To investigate the relationship between a progression of pancreatic cancer and hTERT phosphorylation, we stained each pancreatic precancerous lesions with anti-249T-P antibodies. We found no evidence for anti-249T-P staining in normal ducts, PanIN-1 and −2 but observed robust staining in PanIN-3 **(Figs. 2A**-**2E)**. In pancreatic adenocarcinomas, we found staining of hTERT phosphorylation with anti-249T-P pAbs in the nucleus (**Fig. 2F**). Both normal and atypical multipolar mitotic images exhibited strong hTERT phosphorylation (**Fig. 2G**). Furthermore, some cases displayed strong signals of hTERT phosphorylation in poorly differentiated areas (**Fig. 2H**, upper lesion) and cancer cells that exhibited squamous differentiation (**Fig. 2I**). We examined the number of cells stained with anti-249T-P antibodies, referred as 249T-P positive cells, in each pancreatic ductal lesion (**Fig. 2J**). The number of 249T-P positive cells was the highest in carcinomas, followed by PanIN-3, PanIN-2, PanIN-1, and duct epithelium (*p*<0.05). To consolidate the specificity of the data for IHC with anti-249T-P pAbs, we stained the pancreatic cancer samples with TpMab-1 monoclonal antibody, that also specifically recognize phosphothreonine 249 (**Fig. 1I**), in serial sections prepared from the identical pancreatic cancer lesions (**Figs. S5A**-**S5F**). We observed the identical staining pattern with these antibodies in the nucleus of the same cells and confirmed that phosphorylation signals of hTERT T249 were detected in pancreatic carcinoma. Finally, we found that 249T-P positive cases showed higher incidence of lymph node metastasis than 249T-P negative cases (**Table S2**).

**Fig. 2.**
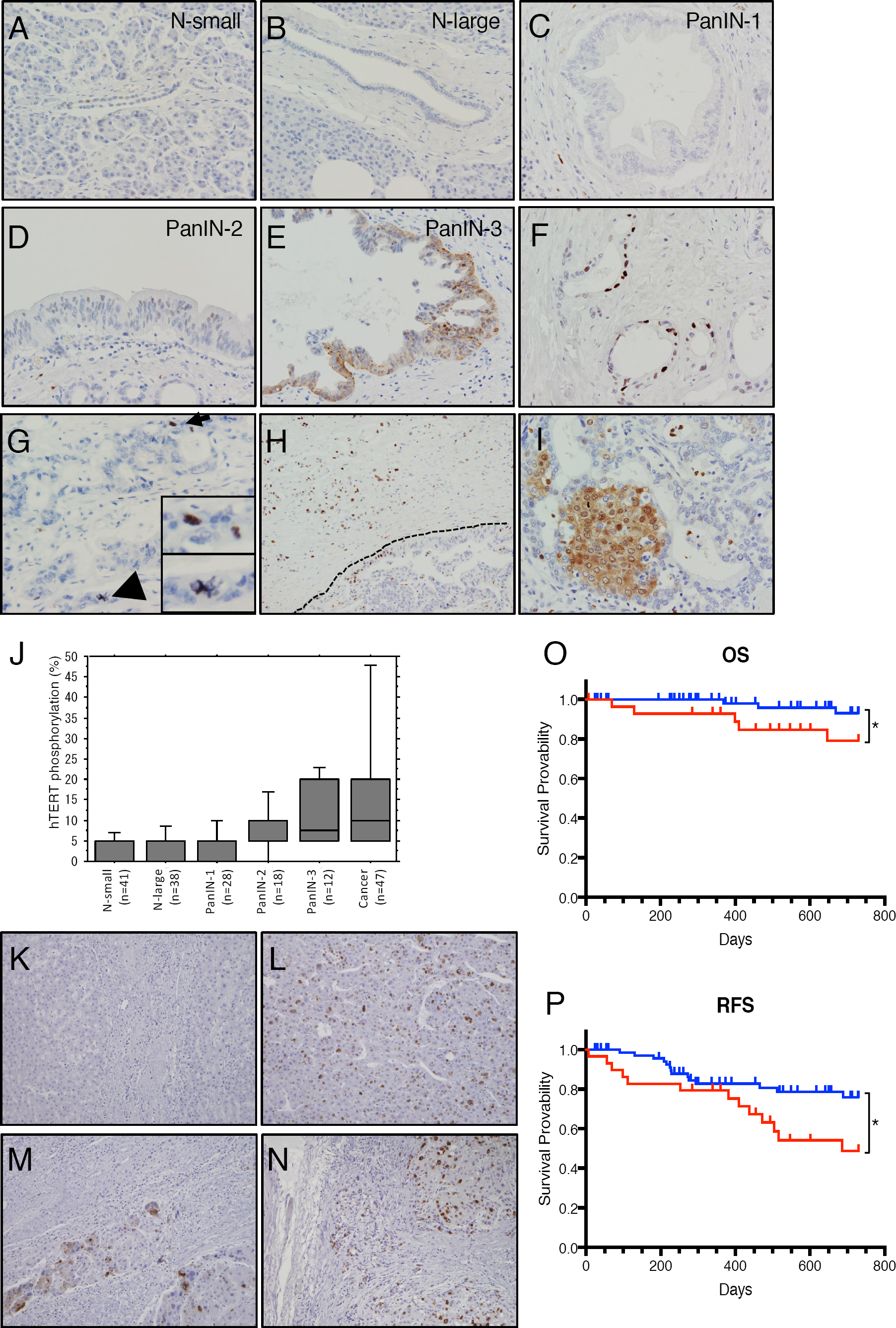
Phosphorylation of hTERT T249 in pancreatic cancer and HCC. **(A-E),** Representative images of precancerous lesions stained with anti-249T-P pAbs. N-small, intercalated duct to intralobular duct; N-large, interlobular duct to main pancreatic duct; PanIN, pancreatic intraepithelial neoplasia; Cancer, pancreatic invasive ductal adenocarcinoma. Original magnification x400. **(F)** Phosphorylation of hTERT T249 was detected in the nucleus of pancreatic adenocarcinomas. Original magnification x400. **(G)** Both normal (arrow) and atypical multipolar mitotic images (arrowhead) exhibited phosphorylation of hTERT T249 (inserts). Original magnification x400. **(H)** Stronger phosphorylation signals of hTERT T249 were displayed in poorly differentiated area (upper lesion) than well differentiated area (lower lesion). Dot line denotes the borderline of “upper lesion” and “lower lesion”. Original magnification x200. **(I)** Cancer cells with squamous differentiation showed strong phosphorylation signals of hTERT T249. Original magnification x400. **(J)** Relationship between pancreatic ductal lesions and hTERT phosphorylation at T249. Box-and-whisker plot of hTERT phosphorylation for each of the pancreatic ductal lesions surgically resected from patients with pancreatic cancer (n=47). *p*<0.05 by Speaman’s rank correlation test. **(K** and **L)** IHC staining with anti-249T-P pAbs in non-cancerous lesions (**K**) and HCC lesions (**L**) obtained from the same patient. Original magnification x200. **(M** and **N)** Strong phosphorylation signals of hTERT T249 were detected in poorly differentiated case (**M**) and in more advanced cancers (**N**). **(O** and **P)** Kaplan-Meier survival analysis of primary patients with HCC grouped for 249T-P positive (red) or negative (blue). Phosphorylation status of hTERT T249 showed statistically significant correlations with overall survival (OS, *p*=0.048, log rank test) (**O**) and relapse-free survival (RFS, *p*=0.0261, log rank test) (**P**) in patients.

In addition, we conducted IHC with anti-249T-P pAbs on hepatocellular carcinoma (HCC) samples. We failed to detect the specific signals in non-cancerous lesions (**Fig. 2K**) but found strong nuclear staining in HCC lesions obtained from the same patient (**Fig. 2L**). Moreover, we found strong hTERT phosphorylation at the T249 in poorly differentiated cases (**Fig. 2M** and **Table S3**) and in more advanced cancers (**Fig. 2N**), suggesting that hTERT phosphorylation at T249 is associated with HCC grade (**Table S3**). In consonance with this observation, we found that patients whose tumors lacked anti-249T-P staining had longer overall survival (OS; *p*=0.048) and recurrence-free survival (RFS; *p*=0.0261) than patients whose cancers showed with anti-249T-P staining (**Figs. 2O** and **2P**). These observations suggest that hTERT phosphorylation at T249 occurs in cancers, more frequently in advanced cancers.

### Consequences of hTERT phosphorylation

To study the consequence of hTERT phosphorylation, we used *λ* phosphatase to remove all phosphate groups from hTERT immunoprecipitated with the anti-hTERT mAb (clone 10E9-2) and then performed a quantitative direct telomerase assay (*15,20*) and an RNA dependent RNA polymerase (RdRP) assay (**Fig. 3A**). We found that phosphatase treatment of hTERT reduced hTERT-dependent RdRP activity but had no measurable effect on telomerase activity.

**Fig. 3.**
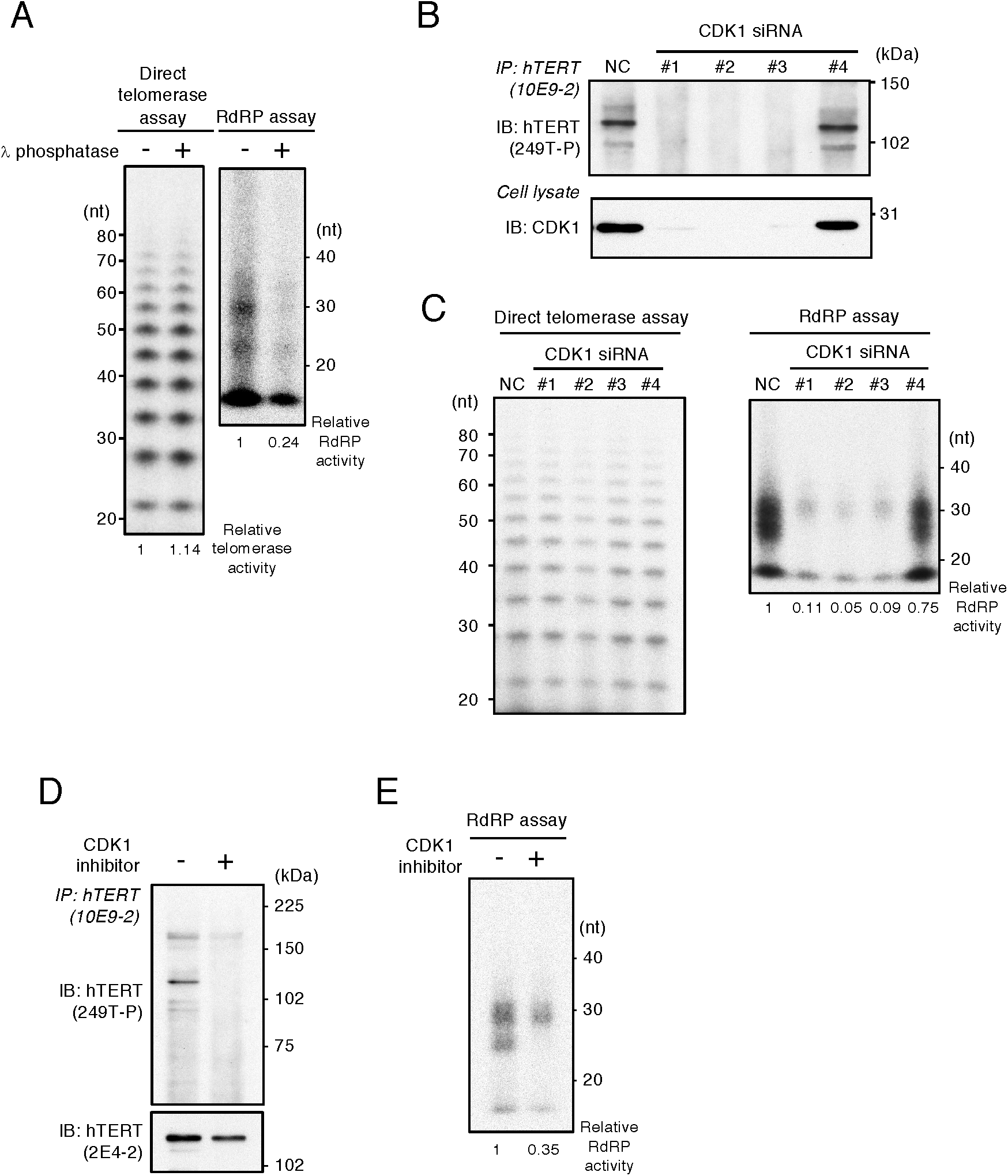
CDK1 phosphorylates hTERT. **(A)** Direct telomerase assay (left panel) and IP-RdRP assay (right panel) using HeLa cells. Immune complexes were treated with *λ* phosphatase to remove the phosphate groups from hTERT proteins. The relative activities of telomerase and RdRP are noted below the panel, respectively. **(B)** Detection of endogenous hTERT proteins using HeLa cells transfected with four different siRNAs specific for *CDK1* followed by nocodazole treatment. Phosphorylation of hTERT proteins were detected by anti-249T-P pAbs (upper panel). Suppression of CDK1 proteins was confirmed by anti-CDK1 mAb (lower panel). **(C)** Direct telomerase assay (left panel) and IP-RdRP assay (right panel) using HeLa cells transfected with four different siRNAs specific for *CDK1*. **(D)** Detection of phosphorylated hTERT proteins using HeLa cells treated with 5 µM of CDK1 inhibitor, RO-3306. The cells were then treated with nocodazole. Phosphorylation of hTERT proteins was detected by anti-249T-P pAbs (upper panel) and hTERT proteins isolated by immunoprecipitation were detected by anti-hTERT mAb (clone 2E4-2) (lower panel). **(E)** IP-RdRP assay using HeLa cells treated with 5 µM of CDK1 inhibitor, RO-3306. The relative RdRP activities are noted below the panel.

To confirm that CDK1 phosphorylates hTERT at T249, we suppressed CDK1 with CDK1-specific siRNAs (**Fig. 3B**). Suppression of CDK1 decreased cell proliferation by 70%, and we observed that introduction of CDK1-specific siRNAs #1, 2, and 3 suppressed hTERT T249 phosphorylation by 98%. Together, these observations strongly suggest that CDK1 is required for the phosphorylation of hTERT at T249.

Moreover, we found that the RdRP activity of hTERT was substantially diminished by CDK1 knockdown while suppression of CDK1 had no effect on telomerase activity (**Fig. 3C**), indicating that phosphorylation by CDK1 is required for the RdRP activity of hTERT. These results suggest that hTERT proteins are phosphorylated by CDK1 in mitosis and the phosphorylation is not required for the telomerase activity but for the RdRP activity of hTERT.

To further confirm that CDK1 specifically phosphorylates hTERT T249, we treated HeLa cells with CDK1 inhibitor, RO-3306 (*21*). Inhibition of CDK1 activity diminished the phosphorylation of hTERT T249 (**Fig. 3D**) and reduced the RdRP activity of hTERT (**Fig. 3E**). Thus, both suppressing CDK1 expression or inhibiting CDK1 kinase activity decreased the phosphorylation of hTERT at T249 and hTERT-mediated RdRP activity.

### Consequences of phosphorylation of hTERT on T249

To further assess the consequences of phosphorylation of hTERT T249 in the human cell lines, we created hTERT mutants where we substituted the threonine 249 of hTERT with alanine (T249A) or glutamic acid (T249E). We introduced these mutants in telomerase-negative human fibroblast BJ cells and analyzed the telomerase activity of the mutant proteins by the telomeric repeat amplification protocol (TRAP) assay. We found that both of these mutant hTERT alleles exhibited telomerase activity equivalent to that observed for wild type hTERT (WT-hTERT) (**Fig. 4A**, left panel). We confirmed the TRAP assay findings by introducing these hTERT mutants in 293T cells and measured the telomerase activity by a quantitative assay as measured by the direct telomerase assay (*15,20*) (**Fig. 4B**). We verified that these mutants retained telomerase activity comparable to WT-hTERT and did not interfere with endogenous hTERT. Consistent with a previous report (*12*), ectopic expression of WT-hTERT protein slightly increased measured telomerase activity. We also assessed whether these two mutants elongated telomeres when expressed in cells by performing Southern blotting for telomere restriction fragments. We analyzed the telomere length in the BJ cells at 8 population doubling (PD) after infection of the two mutants and found that T249A and T249E mutants elongated telomeres (**Fig. 4A**, right panel). These data demonstrated that telomerase activity of the mutants was intact.

**Fig. 4.**
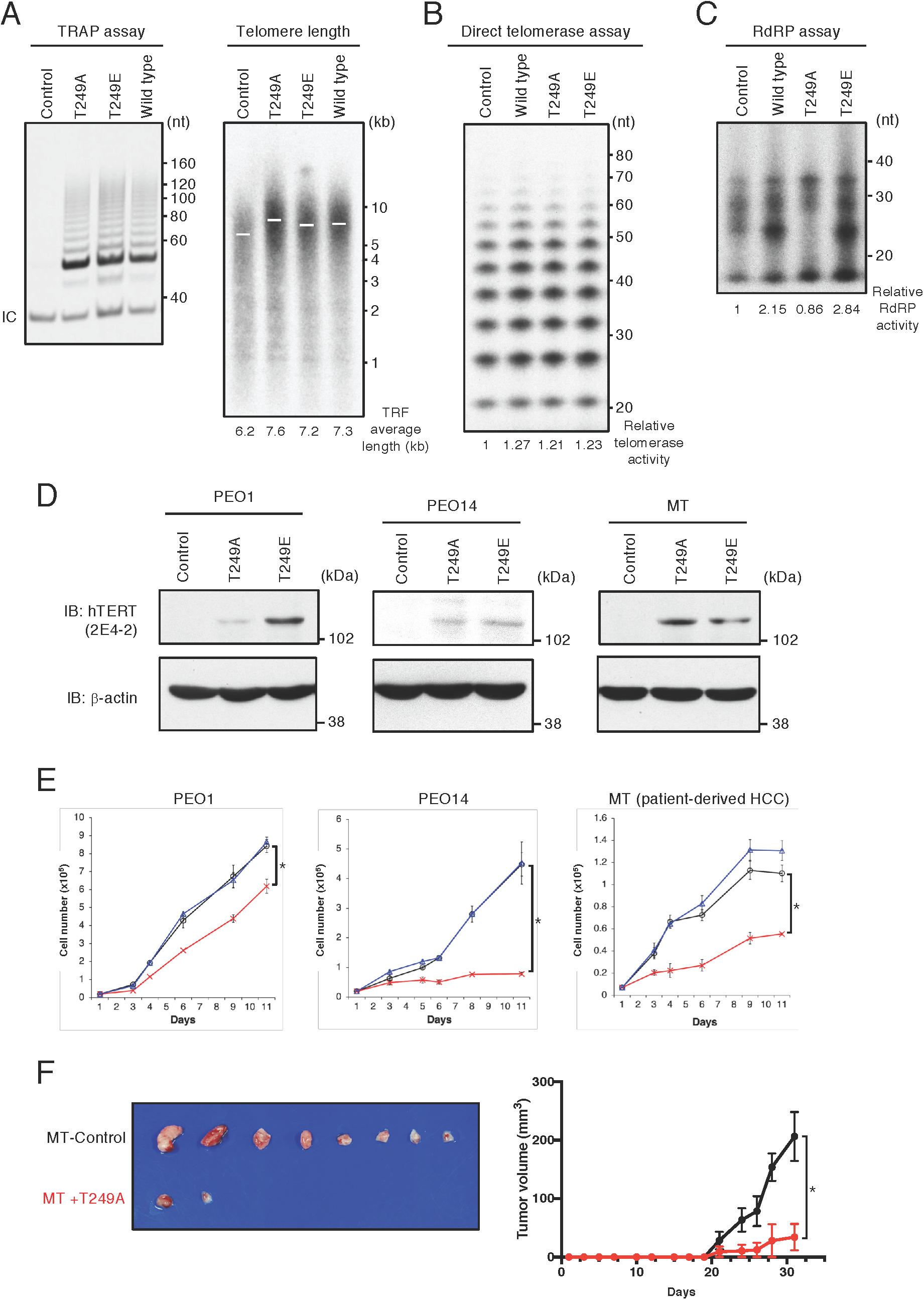
Consequences of hTERT phosphorylation on T249. **(A)** TRAP assay using BJ cells stably introduced with control retrovirus, hTERT mutants or wild type hTERT (left panel). hTERT mutants were created by substitution of the threonine 249 with alanine (T249A) or glutamic acid (T249E). Telomere length in the BJ cells was determined by Southern blot analysis for telomere restriction fragments (right panel). White bars indicate the position of mean radioactivity and the TRF average length were noted below the panel. Telomere structure in BJ cells was analyzed at 8 PD after infection. **(B)** Direct telomerase assay using hTERT proteins immunoprecipitated from 293T cells transfected with control vector, wild type hTERT, T249A or T249E mutants. The relative telomerase activities are noted below the panel. **(C)** IP-RdRP assay using the cells prepared with identical manipulations as used in the panel (**B**). The relative RdRP activities are noted below the panel. **(D)** Immunoblotting to confirm the expression of hTERT mutant proteins in PEO1, PEO14 and MT cells infected with hTERT-T249A, hTERT-T249E or control retroviruses. hTERT proteins were detected by anti-hTERT mAb (clone 2E4-2) (upper panel). β-actin was used as an internal control (lower panel). **(E)** Cell proliferation assay using PEO1, PEO14 and MT (patient-derived HCC) cells infected with hTERT-T249A (red line), hTERT-T249E (blue line) or control retrovirus (black line). This assay was done in triplicate and asterisk indicates statistically significant values (*p*<0.05 by Student’s t-test). **(E)** Expression of transcripts from *hLINE1* in PEO1 and PEO14 cell lines expressing hTERT-T249A or hTERT-T249E mutants as measured by qPCR and normalized with *GAPDH* expression. **(F)** MT cells introduced a control retroviral vector (MT-Control) or hTERT-T249A (MT+T249A) were implanted into eight NOD/SCID mice for each and tumors generated from the cells are presented (left panel). The volume of tumors from mice harboring MT-Control (black line) or MT+T249A (red line) is shown as the mean ± SEM (n=8, *p*=0.0028) (right panel).

We next examined whether introduction of hTERT mutants in 293T cells altered hTERT-dependent RdRP activity (*22*). We found that expression of the phosphorylation-defective mutant hTERT T249A decreased RdRP activity (**Fig. 4C**), while ectopic expression of hTERT T249E increased RdRP activity, suggesting that this mutant acts as a phosphomimetic mutant. These observations confirmed that the phosphorylation of hTERT T249 is required for the RdRP activity but not for reverse transcriptase activity of hTERT.

To further assess the biological effects of phosphorylation of hTERT T249 in cancer cell lines, we expressed hTERT T249A, hTERT T249E, or a control vector expressing only a drug resistance marker into several well-characterized cancer cell lines; two telomerase/RdRP-positive ovarian cancer cell lines (*14*), PEO1 and PEO14, and a telomerase/RdRP-positive patient-derived hepatocellular carcinoma (HCC) MT cells (**Fig. 4D**). In PEO1, PEO14 and MT cells, the expression of hTERT T249A dramatically decreased the cell proliferation (0.73-, 0.18- and 0.50-fold, respectively) while the hTERT T249E showed a minor effect on cell proliferation (1.03-, 1.01- and 1.19-fold, respectively) (**Fig. 4E**). These observations suggest that phosphorylation-defective hTERT T249A interferes with hTERT-mediated RdRP activity and the inhibition of RdRP activity causes inhibitory effects on the cell growth. In contrast, hTERT T249E failed to alter the proliferation of these cell lines.

To further evaluate the influence of hTERT phosphorylation at T249 using patient derived xenograft (PDX) models *in vivo*, we introduced hTERT T249A (MT+T249A) or a control retroviral vector (MT-Control) into MT (patient-derived HCC) cells and implanted these cells subcutaneously into non-obese diabetic/severe combined immunodeficiency (NOD/SCID) mice. Six of eight mice harboring MT+T249A failed to form tumors (**Fig. 4F**, left panel) and the volume of tumors from mice harboring MT+T249A was significantly smaller than those from mice with MT-Control (**Fig. 4F**, right panel; *p*=0.0028). Taken together, these observations indicate that phosphorylation of hTERT at T249 is required for cancer cell proliferation through activation of RdRP activity of hTERT.

### CRISPR-Cas9 genome editing to introduce alanine or glutamic acid substitutions at T249

Although the expression of these hTERT alleles clearly separated hTERT mediated telomerase and RdRP activity, endogenous hTERT is expressed at low levels even in cancer cells. To assess whether the phosphorylation of T249 affected endogenous hTERT function, we used CRISPR-Cas9 genome editing to introduce alanine or glutamic acid substitutions at T249 in the endogenous *hTERT* locus in 293T cells (**Fig. 5A**). After introduction of Cas9 and guide RNAs with donor DNAs into 293T cells, we selected clones and confirmed that the mutations were introduced by PCR amplification, restriction enzyme digestion, and Sanger sequencing (**Figs. 5B** and **5C**). Control cells, Control-CRISPR, were selected from clonal selection of cells expressing Cas9 and no guide RNAs. We confirmed that T249A-CRISPR and T249E-CRISPR were expressed at a level similar to that of endogenous hTERT in the Control-CRISPR cells (**Fig. 5D**, lower panel).

**Fig. 5.**
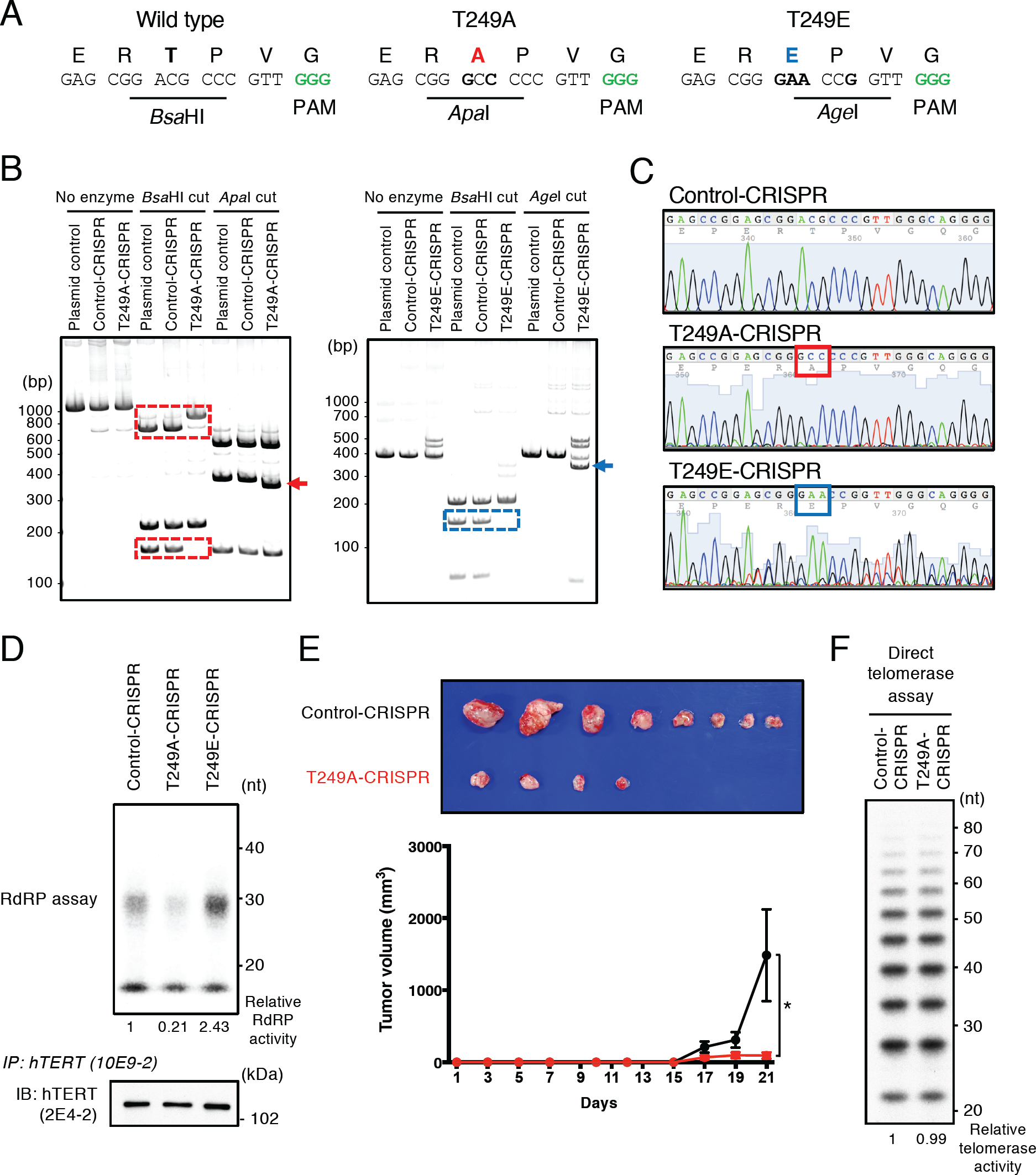
Threonine 249 in the endogenous *hTERT* locus is edited by CRISPR-Cas9 in 293T cells. **(A)** Genome editing strategy to introduce alanine or glutamic acid substitution in the endogenous *hTERT* locus. Green letters denote the PAM (protospacer adjacent motif) sequence for genome editing by CRISPR-Cas9. **(B)** PCR fragments surrounding target regions (1,184 bp for T249A and 456 bp for T249E) were amplified from genomic DNA of genome-edited 293T clones. PCR fragments from pBABE-puro-hTERT (plasmid control), 293T control cells (Control-CRISPR) and mutant cells (T249A-CRISPR, T249E-CRISPR) were digested with the indicated restriction enzymes and electrophoresed in 5% polyacrylamide gels. The bands enclosed with square and indicated by arrows show that the cleavage patterns of PCR fragment were changed in CRISPR-mutant clones. **(C)** Sanger sequencing traces of a control and two mutant clones generated from gel-purified PCR fragments of the genomic DNAs of the respective clones. The sequences boxed in red and blue are the base triplet coding for alanine and glutamic acid in mutant clones, respectively. **(D)** IP-RdRP assay using hTERT proteins immunoprecipitated from 293T CRISPR clones (upper panel). The relative RdRP activities are noted below the panel. hTERT proteins isolated by immunoprecipitation from the cells were detected by anti-hTERT mAb (clone 2E4-2) (lower panel). **(E)** Tumors generated from 293T-Control-CRISPR or 293T-T249A-CRISPR in eight NOD/SCID mice for each are presented (upper panel). The volume of tumors from mice with Control-CRISPR (black line) or T249A-CRISPR (red line) is shown as the mean ± SEM (n=8, *p*=0.0469) (lower panel). **(F)** Direct telomerase assay using hTERT proteins immunoprecipitated from Control-CRISPR or T249A-CRISPR. The relative telomerase activities are noted below the panel.

Using these genome-edited 293T cells, we confirmed that substitution of T249 to alanine ablated hTERT-dependent RdRP activity (**Fig. 5D**, upper panel). Similar to ectopic expression of the mutant proteins (**Fig. 4C**), hTERT with T249E mutation increased the RdRP activity while hTERT with T249A mutation decreased the activity. These results confirmed that phosphorylation of T249 is essential for RdRP activity of hTERT.

To evaluate the direct influence of hTERT phosphorylation at T249 using xenograft models *in vivo*, we inoculated Control-CRISPR or T249A-CRISPR subcutaneously into NOD/SCID mice. Four of eight mice harboring T249A-CRISPR failed to form tumors (**Fig. 5E**, upper panel) and the volume of tumors from mice with T249A-CRISPR was significantly smaller than those from mice with Control-CRISPR (**Fig. 5E**, lower panel; *p*=0.0469). We confirmed that RdRP-deficient hTERT proteins isolated from T249A-CRISPR retained telomerase activity comparable to the hTERT from Control-CRISPR (**Fig. 5F**). Taken together, these observations confirmed that phosphothreonine 249 in hTERT is essential for cancer cell proliferation through activation of RdRP activity of hTERT.

### Effects of hTERT phosphorylation on gene expression

In prior work, we showed that hTERT-mediated RdRP activity affects gene expression by affecting RNA levels (*16*). To identify genes whose expression is altered when hTERT cannot be phosphorylated, we examined gene expression at promoter level using Cap Analysis of Gene Expression (CAGE) method (*23*) for Control-CRISPR and T249A-CRISPR in triplicate. We found 3,482 CAGE peaks, indicating each transcriptional start site (TSS) and the expression level as a promoter activity, that exhibited altered the activity in these cells (FDR<1e-4), where 1,926 and 1,556 were up- and down-regulated in T249A-CRISPR. In particular, we found that three independent CAGE peaks of the transcription factor, forkhead box O4 (*FOXO4*), were upregulated in T249A-CRISPR while the most upstream TSS, p1, showed no change (**Fig. 6A**). We confirmed the increase in expression of FOXO4 proteins in T249A-CRISPR by immunoblotting (**Fig. 6B**). To further confirm that the phosphorylation of hTERT T249 by CDK1 specifically affects the expression of *FOXO4*, we treated HeLa cells with CDK1 inhibitor, RO-3306. Inhibition of CDK1 activity increased the expression of *FOXO4* mRNA in a dose-dependent manner (**Fig. S6**).

**Fig. 6.**
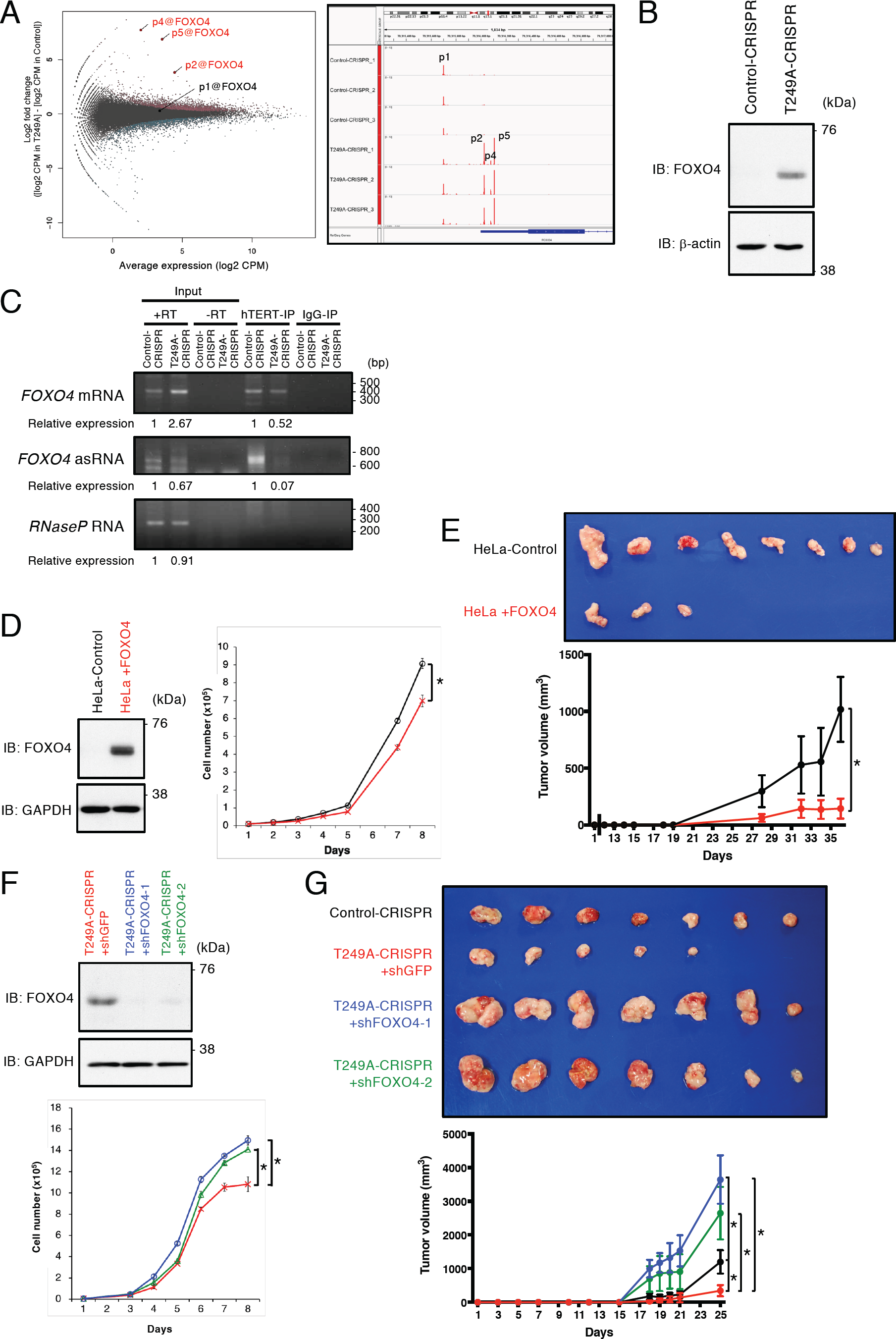
Expression of *FOXO4* is regulated by hTERT phosphorylated on T249. **(A)** MA plot of CAGE peaks differentially expressed between the T249A-CRISPR and Control-CRISPR (left panel). X- and Y-axes indicate average expression levels and fold changes between the two conditions with triplicates, respectively. Individual dots represent CAGE peaks corresponding to promoters defined by the FANTOM5 project (*56*). CAGE peaks with statistical significance are indicated by pink and light blue for higher and lower expression in the T249A-CRISPR, respectively. The CAGE peaks corresponding to *FOXO4* are indicated with red on MA plot and the positions of each *FOXO4* CAGE peak are visualized by Integrative Genomics Viewer (right panel). **(B)** Detection of endogenous FOXO4 proteins in Control-CRISPR and T249A-CRISPR. The proteins were detected by anti-FOXO4 rabbit mAb. *β*-actin was used as an internal control. **(C)** Immune complexes were isolated from Control-CRISPR and T249A-CRISPR with anti-hTERT mAb (clone 10E9-2) and associated RNAs were subjected to RT-PCR. “+RT” and “-RT” indicate the presence and absence of reverse transcriptase, respectively. The relative expressions are noted below the panel. **(D)** Immunoblotting of FOXO4 in HeLa infected with pBABE-hygro (pBh) control and pBh-FOXO4 retroviruses (left panel). Cell proliferation assay using HeLa-Control (black line) or HeLa+FOXO4 (red line) (right panel). The assay was done in triplicate and asterisk indicates statistically significant values (*p*<0.05 by Student’s t-test). **(E)** Tumor appearances generated from subcutaneous injection of HeLa+FOXO4 or HeLa-Control in eight NOD/SCID mice for each are represented (upper panel). The volume curves of tumors derived from mice with HeLa+FOXO4 (red line) or HeLa-Control (black line) are demonstrated as the mean ± SEM (n=8) (lower panel). Asterisk indicates statistically significant values (*p*<0.05 by Student’s t-test). **(F)** Immunoblotting of FOXO4 in T249A-CRISPR expressing FOXO4-specific shRNAs (shFOXO4-1,2) or shGFP control (upper panel). Cell proliferation assay using T249A-CRISPR+shFOXO4-1 (blue line), T249A-CRISPR+shFOXO4-2 (green line) or T249A-CRISPR+shGFP (red line) (lower panel). The assay was done in triplicate and asterisk indicates statistically significant values (*p*<0.05 by Student’s t-test). **(G)** Tumors were established from subcutaneous injection of T249A-CRISPR+shFOXO4-1, −2, T249A-CRISPR+shGFP or Control-CRISPR in seven NOD/SCID mice for each (upper panel). Tumor-volume curves of xenografts derived from T249A-CRISPR+shFOXO4-1 (blue line), T249A-CRISPR+shFOXO4-2 (green line), T249A-CRISPR+shGFP (red line), or Control-CRISPR (black line) are shown as the mean ± SEM (n=7) (lower panel). Asterisk indicates statistically significant values (*p*<0.05 by Student’s t-test).

Gene ontology (GO) term analysis showed enrichment of genes associated with “cell cycle phase transition” (GO:0044770, **Table S4**) within the upregulated genes in the T249A-CRISPR. Notably, *FOXO4* is annotated in this GO term and functions as tumor suppressor proteins in various cancers by preventing proper cell cycle regulation (*24-29*). These findings indicate that phosphorylation of T249 on hTERT regulates the expression of *FOXO4* negatively in cancer cell lines and increase of *FOXO4* expression in T249A-CRISPR is correlated with a decrease of tumor formation.

To determine whether *FOXO4* is a direct target of hTERT phosphorylated at T249, we performed IP-RT-PCR experiments with RNAs isolated from hTERT-IP samples of Control-CRISPR and T249A-CRISPR. We isolated hTERT-RNA complexes by immunoprecipitation with an anti-hTERT mAb (clone 10E9-2) from Control-CRISPR and T249A-CRISPR cells, and purified RNAs from the immune complexes with DNase treatment. cDNAs were synthesized with strand-specific primers, and we performed PCR using primers to amplify the first exon of *FOXO4* mRNAs. We confirmed that endogenous hTERT immunoprecipitated from Control-CRISPR interacts with the first exon of *FOXO4* mRNAs and that there was a 50% reduction of hTERT associated with *FOXO4* mRNAs isolated from T249A-CRISPR cells (**Fig. 6C**). We also found that hTERT from Control-CRISPR interacts with antisense RNAs (asRNAs) of *FOXO4* mRNA under condition in which we failed to recover *RNase P* RNA (*22*).

Taken together, we propose a model for regulation of *FOXO4* expression via phosphorylation of hTERT at T249 (**Fig. S7**). Specifically, in Control-CRISPR, hTERT proteins phosphorylated at T249 bind to the first exon of *FOXO4* mRNAs. These RNAs serve as templates for double-stranded RNA (dsRNA) synthesis by hTERT-RdRP. These dsRNA complexes are degraded and the expression of *FOXO4* is reduced. In T249A-CRISPR, RdRP-deficient hTERT-T249A proteins interact with less *FOXO4* mRNAs than hTERT proteins in Control-CRISPR. Without producing asRNA, more *FOXO4* mRNAs are retained in the T249A-CRISPR and increase of *FOXO4* expression causes tumor suppression via cancer cell growth arrest.

To corroborate the effects of *FOXO4* expression in cancer cells, we established stable cell lines introduced with pBABE-hygro (pBh) control vector or *FOXO4*-expressing retroviral vector (pBh-FOXO4) in HeLa cells. Increase of FOXO4 protein expression introduced with pBh-FOXO4 (HeLa+FOXO4) was confirmed by immunoblotting whereas expression level of FOXO4 introduced with pBh (HeLa-Control) was below the detection limit of anti-FOXO4 antibody (**Fig. 6D**, left panel). Up-regulation of FOXO4 significantly inhibited the cell proliferation *in vitro* by 77.0% (**Fig. 6D**, right panel). To evaluate the effects of FOXO4 overexpression in HeLa using xenograft models, we inoculated HeLa-Control or HeLa+FOXO4 subcutaneously into NOD/SCID mice. Retroviral expression of FOXO4 resulted in reduction of tumor formation in five of eight mice (**Fig. 6E**, upper panel). The volume of tumors from mice harboring HeLa+FOXO4 was significantly smaller than those from mice with HeLa-Control (**Fig. 6E**, lower panel, *p*=0.0092). Taken together, these observations confirmed that increase of *FOXO4* expression in T249A-CRISPR brings about a decrease of tumor formation *in vivo*.

To further assess the biological influences of increase of *FOXO4* expression in T249A-CRISPR, we established stable cell lines expressing short hairpin RNAs (shRNAs) specific for *FOXO4* (shFOXO4-1, 2) or shGFP control in T249A-CRISPR. Knockdown of FOXO4 protein expression in the cell lines was confirmed by immunoblotting (**Fig. 6F**, upper panel). Down-regulation of *FOXO4* in T249A-CRISPR significantly increased the cell proliferation by 1.31- and 1.26-fold, respectively (**Fig. 6F**, lower panel). To evaluate the effects of *FOXO4* suppression in T249A-CRISPR on tumor formation, we used xenograft models by the subcutaneous inoculation of T249A-CRISPR expressing shFOXO4 (T249A-CRISPR+shFOXO4-1, 2) or shGFP (T249A-CRISPR+shGFP) into NOD/SCID mice. While five of seven mice with T249A-CRISPR+shGFP formed xenograft tumors, T249A-CRISPR+shFOXO4-1 and −2 developed tumors in all seven mice (**Fig. 6G**, upper panel). Compared to T249A-CRISPR+shGFP and Control-CRISPR, T249A-CRISPR+shFOXO4 exhibited a higher tumor growth rate and the volume of tumors from mice with T249A-CRISPR+shFOXO4-1 and −2 was significantly larger than those from mice with T249A-CRISPR+shGFP (**Fig. 6G**, lower panel; *p*=0.0008 and *p*=0.0136, respectively). These observations verified that perturbation of *FOXO4* expression by specific shRNAs rescued the defects in tumor formation ability in T249A-CRISPR.

## Discussion

Telomerase plays an essential role in maintaining telomeres. The expression of the catalytic subunit of telomerase hTERT is upregulated in the majority of human cancers, most often due to mutations in its promoter. Here we demonstrate that hTERT is phosphorylated by CDK1 in mitosis and that this phosphorylation event affects a second hTERT function, RNA dependent RNA polymerase activity without affecting hTERT function at telomeres. This mitotic function of hTERT correlates with tumor aggressiveness and is necessary for tumor formation. Together these observations separate the telomere and non-telomere directed functions of hTERT and identify a novel mechanism that regulates hTERT RdRP activity.

It is well accepted that telomerase recruitment to telomeres occurs in S-phase in many organisms (*10,30*). While these observations indicate that telomerase recruitment to telomeres and telomerase activation are regulated in a cell cycle dependent manner, there have been two reports that studied telomerase activity outside the S-phase (*31,32*). Specifically, these reports showed that *Xenopus* telomerase activity was active in mitotic phase and modulated chromatin configuration. In consonance with these findings, we previously reported that hTERT localizes to both mitotic spindles and centromeres, that hTERT protein expression level is enriched in mitosis, and that hTERT-RdRP activity is present in nuclear extracts derived from several cancer cell lines arrested in mitosis (*14,16,33*). Here we have confirmed and extended these findings by identifying the mitotic kinase CDK1 as a key regulator of hTERT RdRP but not telomerase activity.

Several laboratories have shown that hTERT is phosphorylated (*17,34-36*). Specifically, four phosphorylation sites have been reported, serine 227 (*17*), serine 457 (*36*), tyrosine 707 (*35*) and serine 824 (*34*). Serine 227 is located in a canonical nuclear localization signal, is phosphorylated by Akt1, and promotes nuclear localization of hTERT (*17*). Serine 824 is also phosphorylated by Akt1 and is necessary for hTERT-mediated telomerase activity (*34*). Serine 457 is phosphorylated by DYRK2 during G2/M and promotes ubiquitination of hTERT proteins (*36*). Tyrosine 707 is phosphorylated by C-Src under oxidative stress to alter subcellular localization of hTERT (*35*). We were unable to detect these sites by MS analysis in this study with mitotic cells. These phosphorylation sites were identified from consensus sequences for specific kinases, and MS analysis was not conducted in previous reports. We speculate that the amount and level of phosphorylation in these sites may be difficult to detect using MS analysis. In addition, previous reports demonstrated that phosphorylation affect either telomerase activity or subcellular localization of hTERT whereas this study identifies the novel phosphorylated site that does not affect telomerase activity.

Threonine 249 in hTERT is conserved in human and primates but not in the other mammals and yeast (**Fig. 1E**), suggesting that phosphorylation-dependent RdRP activity of TERT is restricted to primates. When we introduced substitutions of T249 (T249A and T249E) into endogenous hTERT, we found that these mutants did not affect telomerase activity at telomeres but instead affected RdRP activity and inhibited the tumorigenic potential of human cancer cell lines. These observations suggest that phosphorylation of hTERT at T249 contributes to tumor formation independent of its reverse transcriptase activity to elongate telomeres. Furthermore, we found that phosphorylation of hTERT at T249 regulates the expression of the tumor suppressor gene, *FOXO4*, in cancer cell lines (**Fig. S7**). Expression levels of *FOXO4* inversely correlate with tumor formation and incidence of clinical metastasis (*24-29*). Consistent with *FOXO4* providing a role of tumor suppressor, T249A-CRISPR expressing higher levels of *FOXO4* formed less and smaller tumors in mouse xenograft model (**Figs. 5E** and **6G**). Our findings indicate that phosphorylation of hTERT T249 regulates *FOXO4* expression negatively by the RdRP activity in cancer cells and is implicated in pivotal event for carcinogenesis.

In addition, we have identified CDK1 as a kinase that phosphorylates hTERT at T249 in mitosis. CDK1 is essential for cell division in the embryo and deficiency in CDK1 in mouse model results in embryonic lethality in the first cell divisions (*37*). Moreover, CDK1 is sufficient to drive the cell cycle in all cell types (*37*). Previous reports demonstrated that CDK1 expression is upregulated in many human malignant tumor tissues and that CDK1 activity correlates with the prognosis of patients with tumors (*38-43*). In addition, loss of CDK1 in the liver in CDK1 conditional knockout mice showed complete resistance against tumorigenesis (*44*) indicating that CDK1 is required for tumorigenesis in liver cancer. Here we report a link between CDK1 and phosphorylation of hTERT at T249, and the phosphorylation occurs more frequently in aggressive and advanced cancers in clinical samples, suggesting an additional role for CDK1 in cancer progression. These observations implicate CDK1 and phosphorylation of hTERT T249 as an approach to inhibit a key cancer-associated function of hTERT.

## Materials and Methods

### Antibodies

Anti-hTERT mouse monoclonal antibodies (mAbs, clones 10E9-2 and 2E4-2) were generated and described the specificity as reported previously (*16*). Anti-hTERT mouse mAb (clone 2E4-2), anti-hTERT rabbit mAb (Abcam, ab32020), anti-phospho-histone H3 (Ser10) rabbit polyclonal antibodies (pAbs) (Merck, 06-570), anti-cdc2 (POH1) mouse mAb (Cell Signaling Technology, 9116), anti-*β*-actin (AC-15) mouse mAb (Sigma-Aldrich, A5441), anti-FOXO4 rabbit mAb (Abcam, ab128908) and anti-GAPDH mouse mAb (MBL Co., Ltd., M171-3) were used for immunoblotting (IB). Anti-hTERT mAb (clone 10E9-2: MBL Co., Ltd., M216-3) and anti-hTERT sheep pAbs (Abbexa Ltd., abx120550) were used for immunoprecipitation (IP).

### Generation of phospho-specific polyclonal antibodies

To generate phosphospecific antibodies against phosphorylated threonine 249 of hTERT (anti-249T-P), hTERT phosphopeptide ^244^CEPERpTPVGQG^254^ was used to immunize rabbits. The antibodies were purified from the antisera by using hTERT phosphopeptide-conjugated Resin and then further purified by passing it through hTERT nonphosphopeptide-conjugated Resin.

### Generation of hybridoma producing phospho-specific monoclonal antibody

Female 4-week-old BALB/c mice were purchased from CLEA Japan and kept under specific pathogen-free conditions. The Animal Care and Use Committee of Tohoku University approved animal experiments to produce antibody in this study. BALB/c mice were immunized by intraperitoneal (i.p.) injection of 100 µg of hTERT phosphopeptide ^244^CEPERpTPVGQG^254^ together with Imject Alum (Thermo Fisher Scientific). After two additional immunizations of 100 µg, a booster injection of 100 µg was given i.p. 2 days before spleen cells were harvested. The spleen cells were fused with P3U1 cells (ATCC) using PEG1500 (Roche Diagnostics,). The hybridomas were grown in RPMI 1640 medium (Nacalai Tesque) at 37°C in a humidified atmosphere containing 5% CO2 and 95% air, supplemented with 10% heat-inactivated fetal bovine serum (Thermo Fisher Scientific), hypoxanthine, aminopterin, and thymidine (HAT) selection medium supplement (Thermo Fisher Scientific), and 5% BriClone Hybridoma Cloning Medium (QED Bioscience). One hundred units/ml penicillin, 100 µg/ml streptomycin, and 25 µg/ml amphotericin B (Nacalai Tesque) were added to the culture medium. Plasmocin (5 µg/ml; InvivoGen) was also used to prevent *Mycoplasma* contamination. The culture supernatants were screened using enzyme-linked immunosorbent assay (ELISA) for binding to hTERT phosphopeptide and nonphosphopeptide.

### Enzyme-Linked Immunosorbent Assay (ELISA)

Peptides were immobilized on Nunc Maxisorp 96-well immunoplates (Thermo Fisher Scientific) at 1 µg/ml. After blocking with SuperBlock T20 (PBS) Blocking Buffer (Thermo Fisher Scientific), the plates were incubated with culture supernatant with subsequent 1:2000 diluted peroxidase-conjugated anti-mouse IgG (Agilent Technologies). The enzymatic reaction was conducted with 1-Step Ultra TMB-ELISA (Thermo Fisher Scientific). The optical density was measured at 655 nm using an iMark microplate reader (Bio-Rad Laboratories). These reactions were performed with a volume of 50-100 µl at 37°C.

### Cell culture and mitotic cell synchronization

The human cervical carcinoma cell line HeLa, the SV40-transformed human embryonic kidney cell line HEK-293T (293T) and human patient-derived hepatocellular carcinoma cell line MT were cultured in DMEM supplemented with 10% heat-inactivated fetal bovine serum (IFS). The human ovarian carcinoma cell lines PEO1 and PEO14 were cultured in RPMI-1640 medium supplemented with 10% IFS and 2 mM sodium pyruvate (Gibco). The human foreskin fibroblast cell line BJ was cultured in K/O DMEM/Medium 199 (4:1) supplemented with 15% IFS and L-Gultamine (Nacalai Tesque).

Mitotic cell synchronization was performed as described previously (*16*). Briefly, cells were switched to medium containing 2.5 mM thymidine (Nacalai Tesque) and incubated for 24 hours. Six hours after release, cells were incubated in medium containing 0.1 µg/ml nocodazole (Sigma-Aldrich) for 16 hours. After shake-off, mitotic cells were retrieved. Cells arrested in mitosis with nocodazole were confirmed by IB using anti-phospho-histone H3 (Ser10) antibodies.

### Transfection of siRNAs

For suppression of hTERT or CDK1 expression, HeLa cells were transfected with siRNAs using Lipofectamine 2000 (Thermo Fisher Scientific). After 48 h of incubation, cells were treated with 0.1 µg/ml nocodazole for 16 hours. The sequences of siRNAs against hTERT (*13*) are listed in **Table S5**. MISSION siRNAs Hs_CDC2_4049_s and Hs_CDC2_4049_as, Hs_CDC2_4053_s and Hs_CDC2_4053_as, Hs_CDC2_6210_s and Hs_CDC2_6210_as, and Hs_CDC2_4050_s and Hs_CDC2_4050_as were used for CDK1 siRNA #1, CDK1 siRNA #2, CDK1 siRNA #3 and CDK1 siRNA #4, respectively. MISSION siRNA Universal Negative Control #1 (Sigma-Aldrich) was used as a negative control.

### IP-IB of hTERT

The IP-IB assay was performed as described previously (*16*). In brief, 1× 10^7^ cells were lysed in 1 ml of Lysis buffer A (0.5% NP-40, 20 mM Tris-HCl (pH 7.4), and 150 mM NaCl). After sonication, lysates were cleared of insoluble material by centrifugation at 21,000 × *g* at 4°C for 15 min. One milliliter of lysate was pre-absorbed with 40 µl of Pierce Protein A Plus Agarose (Thermo Fisher Scientific) for 30 min at 4°C. Pre-absorbed lysate was mixed with 10 µg of anti-hTERT mAb (clone 10E9-2) or 30 µg of anti-hTERT pAbs (abx120550) and 40 µl of Pierce Protein A Plus Agarose, and incubated overnight at 4°C. Immune complexes were washed three times with Lysis buffer A and eluted in 2× SDS loading buffer (2% *β*-mercaptoethanol, 20% glycerol, 4% SDS, and 100 mM Tris-HCl (pH 6.8)), and then subjected to SDS-PAGE in 8% polyacrylamide gels. Anti-hTERT mouse mAb (clone 2E4-2) and Mouse TrueBlot ULTRA Anti-Mouse Ig HRP (Rockland) or anti-hTERT rabbit mAb (ab32020) and anti-rabbit IgG HRP (GE Healthcare) were used for IB to detect whole hTERT proteins. Anti-phospho hTERT rabbit pAbs (anti-249T-P) and anti-rabbit IgG HRP (GE Healthcare) or anti-phospho hTERT mouse mAb (TpMab-1) and anti-mouse IgG HRP (GE Healthcare) were used for IB to detect phosphorylated hTERT proteins.

For the *λ* phosphatase treatment, the beads suspension with immune complexes was treated with 2,000 U of *λ* protein phosphatase (Bio Academia) and 2 mM MnCl2 in *λ*-PPase reaction buffer (50 mM Tris-HCl (pH 7.6), 100 mM NaCl, 2 mM DTT, 100 µM EDTA and 0.01% Brij 35) and incubated at 30°C for 30 min.

### Identification of phosphopeptides by mass spectrometric protein profiling

The protein samples immunoprecipitated with anti-hTERT mAb (clone 10E9-2) were separated by 12.5% SDS-PAGE and subjected to in-gel digestion using trypsin (*45*). The tryptic digests were subjected to liquid chromatography coupled with nanoelectrospray tandem mass spectrometry (Finnigan LTQ Orbitrap XL mass spectrometer; Thermo Fisher Scientific). The Mascot software package (version 2.5.1; Matrix Science) was used to search for the mass of each peptide ion peak against the SWISS-PROT database (Homo sapiens, 20,205 sequences in the Swiss prot_2015_09.fasta file) using the following parameters: 1 missed cleavage; fixed modification: carboxymethylation (C); variable modification: oxidation (M), phosphorylation (ST), phosphorylation (Y); search mode: MS/MS ion search with decoy database search included; peptide mass tolerance ± 10 ppm; MS/MS mass tolerance ± 0.8 Da; peptide charge: 2+ and 3+. All peptide information of TERT proteins detected in this study is shown in **Supplementary Data S1**.

### *In vitro* kinase assay

For recombinant protein expression using cell-free synthesis-coupled transcription–translation (*46*), the target cDNA fragment, corresponding to 191-306 residues of hTERT was sub-cloned into the pCR2.1-TOPO vector (Thermo Fisher Scientific). Proteins, subjected to affinity purification, were N-terminally fused with a modified natural poly-histidine N11-tag (amino-acid sequence: MKDHLIHNHHKHEHAHAEH) with a TEV (Tobacco Etch virus) protease recognition site and a GSSGSSG linker sequence. These sequences were introduced using TOPO cloning. The cell-free synthesized N11-tagged hTERT_191-306 protein was purified using an AKTA 10S system (GE Healthcare) with a HisTrap columns (GE Healthcare); the AKTA 10S system was washed with a concentration gradient buffer (50 mM Tris-HCl buffer at pH 8.0, containing 1 M NaCl and 10 mM imidazole). The N11-tagged recombinant proteins were eluted with a concentration gradient of imidazole (from 10 to 500 mM) in elution buffer (50 mM Tris-HCl buffer at pH 8.0, containing 0.5 M NaCl). Imidazole was removed by overnight dialysis at 4°C in wash buffer. The affinity-tags were removed by incubation at 4°C for 20 hours with TEV protease. The resulting tag-cleaved proteins were purified by ion exchange chromatography with a HiTrap SP column (GE Healthcare) and gel filtration with the final buffer (25 mM Tris-HCl buffer at pH 7.0, containing 450 mM NaCl, 0.25mM TCEP) using HiLoad 16/600 Superdex columns (GE Healthcare).

For baculovirus–insect cell expression of active form of human IKK2 (IKK2_2-664), we used the Bac-to-Bac Baculovirus Expression System (Thermo Fisher Scientific). The target cDNA fragment, corresponding to 2-664 residues of human IKK2, was sub-cloned into the pDEST vector (Thermo Fisher Scientific). Polyhistidine affinity-tagged IKK2_2-664 was expressed in insect Sf9 cells at a multiplicity of infection (MOI) of 1.0 with a recombinant baculovirus that expresses IKK2_2-664. The cells from each culture were harvested 48 hours post-infection, and the cell pellets were washed once with PBS and were immediately frozen in liquid N2. The recombinant IKK2_2-664 protein was purified as described previously (*47*).

Purified hTERT_191-306 proteins were incubated with CDK1-cyclin B (New England Biolabs) or purified IKK2_2-664 proteins. *In vitro* kinase assays were carried out in 30 µl of reaction buffer (50 mM Tris-HCl pH7.5, 10 mM MgCl2, 0.1 mM EDTA, 2 mM DTT, 0.01% Brij 35) containing 4 mM ATP, 10 µM of hTERT_191-306 proteins and 30 units of CDK1-cyclin B or 1 µM of IKK2_2-664. Reactions were incubated at 37°C for 2 hours. Reaction samples were terminated by adding 2.5x SDS sample loading buffer (5% (w/v) SDS, 250 mM DTT, 15% (v/v) glycerol, 140 mM Tris–HCl pH 6.8, and 0.01% (w/v) bromophenol blue), boiled and subjected to Phos-tag SDS– PAGE (*19*).

For mass spectrometry analyses, gel regions, containing proteins, were excised and digested with trypsin (Promega) for 20 hours at 37°C. The resulting peptides were analyzed by LC-ESI-MS/MS (liquid chromatography–electrospray ionization tandem mass spectrometry) at the Support Unit for Bio-Material Analysis in RIKEN CBS Research Resources Center. All peptide information detected in this study is shown in **Table S1**.

### IP-RdRP assay

The IP-RdRP assay was performed as described previously (*16*). TERT protein was immunoprecipitated from human cell lines as described for the IP-IB assay with anti-hTERT mAb (clone 10E9-2). The bead suspension with immune complexes was washed four times with 1× acetate buffer (10 mM HEPES-KOH (pH 7.8), 100 mM potassium acetate, and 4 mM MgCl2) containing 10% glycerol, 0.1% Triton-X, and 0.06× cOmplete EDTA-free (Roche), and once with AGC solution (1× acetate buffer containing 10% glycerol and 0.02% CHAPS) containing 2 mM CaCl2. The bead suspension was treated with 0.25 unit/µl *Micrococcal Nuclease* (Takara Bio) at 25°C for 15 min. Immunoprecipitates were subsequently washed twice with AGC solution containing 3 mM EGTA and once with 1× acetate buffer containing 0.02% CHAPS. Forty microliter of reaction mixture was prepared by combining 20 µl of the bead suspension with 6 µl of [*α*-^32^P] UTP (3,000 Ci/mmol) and 25 ng/µl (final concentration) of RNA template, and incubated at 32°C for 2 hours. The sequence of RNA template is as follows: 5’-GGGAUCAUGUGGGUCCUAUUACAUUUUAAACCCA-3’ (*48*). This RNA has hydroxyl groups at both the 5’ and 3’ ends. The final concentrations of ribonucleotides were 1 mM ATP, 0.2 mM GTP, 10.5 µM UTP, and 0.2 mM CTP. The resulting products were treated with Proteinase K to stop the reaction, purified several times with phenol/chloroform until the white interface disappeared, and precipitated using ethanol. The RdRP products were treated with RNase I (2 U, Promega) at 37°C for 2 hours to digest single-stranded RNAs completely, followed by Proteinase K treatment, phenol/chloroform purification, and ethanol precipitation. The products were electrophoresed in a 10% polyacrylamide gel containing 7 M urea, and detected by autoradiography.

### Generation of phosphorylation-inhibitory or phosphomimetic mutant plasmids

For site-directed mutagenesis, QuikChange II XL site-directed mutagenesis Kit (Agilent Technologies) was used. In brief, site-directed mutagenesis PCRs were performed using PfuUltra High Fidelity DNA polymerase following the manufacturer’s protocol with either the pBABE-puro-hTERT retroviral vector or the pNK-FLAG-Z-hTERT expression vector as template plasmids. Sequences of mutagenic primers are listed in **Table S5**. PCR products were digested with *Dpn*I restriction enzyme for 1 hour at 37°C and then transformed into XL-10-Gold ultracompetent cells. Mutations were confirmed by Sanger sequencing.

### Telomere analysis

For the telomeric repeat amplification protocol (TRAP) assay, 1× 10^5^ cells were suspended in 200 µl of TRAP lysis buffer (10 mM Tris-HCl (pH 7.5), 1 mM MgCl2, 1 mM EGTA, 0.5% CHAPS, 10% glycerol, 100 µM Pefabloc SC, and 0.035% 2-mercaptoethanol). The TRAP assay was performed with 5 µl of the suspension as described previously (*13*).

To measure telomere length by Southern blotting, genomic DNAs were isolated, digested with *Hin*fI and *Afa*I, electrophoresed in a 0.8% agarose gel and hybridized with a ^32^P-labeled telomeric (CCCTAA)3 probe as described previously (*49*).

### Direct telomerase assay

For the direct telomerase assay, we modified the original methods (*15,20*). Briefly, TERT protein was immunoprecipitated from human cell lines as described for the IP-IB assay without sonication. Immune complexes were washed three times with Lysis buffer A, and then suspended in 30 µl of TRAP lysis buffer. The direct telomerase assay was carried out with 10 µl of the suspension and 40 µl of reaction mixture (2.5 µl of [*α*-^32^P] dGTP (6,000 Ci/mmol), 50 mM Tris-HCl (pH8.0), 50 mM KCl, 1 mM MgCl2, 1.25 mM Spermidine, 5 mM 2-mercaptoethanol, 2.5 mM dTTP, 2.5 mM dATP, 25 µM dGTP and 2.5 µM a5 primer (5’-TTAGGGTTAGGGTTAGCGTTA-3’)) by incubation at 37°C for 2 hours. The resulting products were purified with phenol/chloroform, and precipitated using ethanol. The products were electrophoresed in a 10% polyacrylamide gel containing 7 M urea, and detected by autoradiography.

### Generation of stable cell lines and proliferation assay

Amphotropic retroviruses were created as described (*50*) using the retroviral vectors pBABE-puro, pBABE-puro-hTERT T249A or pBABE-puro-hTERT T249E for making stable cell lines expressing hTERT mutants. After infection, polyclonal cell populations were purified by selection with puromycin (2 µg/ml) for 3 days.

We used the following short hairpin RNA (shRNA) vectors, shFOXO4-1 (TRCN0000010291) and shFOXO4-2 (TRCN0000039720), constructed by The RNAi Consortium and the sequences are listed in **Table S5**. shGFP was used as the control. These vectors were used to make amphotropic lentiviruses, and the cell populations were selected with puromycin (2 µg/ml) for 3 days.

For generating stable cell lines expressing FOXO4, we constructed the retroviral vector pBABE-hygro-FOXO4. After infection of pBABE-hygro or pBABE-hygro-FOXO4, the cells were selected with hygromycin B (200 µg/ml) for 7 days.

To generate proliferation curves, cells were plated in triplicate and counted in a Z2 Particle Count and Size Analyzer (Beckman-Coulter).

### RT-PCR and quantitative RT-PCR (qPCR)

Total cellular RNA was isolated using TRIzol (Thermo Fisher Scientific), treated with RQ1 DNase (Promega), and subjected to RT-PCR and qPCR. For IP-RT-PCR, RNA samples were extracted from the immune complexes as described for the IP-IB assay without sonication. Immune complexes were washed three times with Lysis buffer A containing 300 mM NaCl and associated RNAs were isolated with TRIzol. The RT reaction was performed with oligo(dT)12-18 (Thermo Fisher Scientific) primer or target- and strand-specific primers using PrimeScript Reverse Transcriptase (Takara Bio) for 60 min at 42°C, followed immediately by PCR. qPCR was performed with a LightCycler 480 II (Roche) using LightCycler 480 SYBR Green I Master (Roche) according to the manufacture’s protocols. Sequences of PCR primers are listed in **Table S5**.

### Genome editing using CRISPR-Cas9

The PuroCas9 plasmid which consists of three cassettes for Cas9 cDNA, chimeric guide RNA and puromycin cDNA was generated by modifying the px330 plasmid (addgene #42230) (*51*). Cas9 and puromycin resistance genes are expressed under the control of the CAG promoter (*52*) and the PGK promoter (*53*), respectively. The guide RNA sequence targeted against hTERT was cloned into the *Bbs*I site of the PuroCas9 plasmid (PuroCas9-hTERT). For generating T249A-CRISPR and T249E-CRISPR, the T249A donor plasmid or T249E single-stranded oligo included the point mutation leading to amino acid change were transfected with PuroCas9-hTERT using FuGeneHD (Promega) into 293T cells, respectively. For generating Control-CRISPR, PuroCas9 plasmid was transfected using FuGeneHD into 293T cells. Following selection with puromycin, single cell was cloned. Genomic DNA was extracted with the GenElute Mammalian genomic DNA miniprep kit (Sigma) according to the manufacturer’s instructions. Presence of mutation in single-cell clones was confirmed by PCR amplification, restriction enzyme digestion and Sanger sequencing. Sequences of PCR primers, donor plasmid/oligo and guide RNA are listed in **Table S5**.

### Xenotransplantation

Nonobese diabetic/severe combined immunodeficiency (NOD/SCID) mice (Charles River Laboratories, Inc.) were used as recipients for xenotransplantation. 1x 10^5^ cells of MT cells introduced retroviral vectors or 293T-CRISPR cells were suspended in a mixture of serum-free medium and Matrigel (BD Biosciences; 1:1 volume). The mixture was injected subcutaneously through a 26-gauge needle into the right dorsal areas of anesthetized NOD/SCID mice. We monitored tumor formation and tumor size every two or three days, and dissected out the tumors within a month after engraftment.

### CAGE sequencing and data analysis

Of each of the total RNAs extracted from two triplicates (Control-CRISPR and T249A-CRISPR), 3 µg was used to prepare a sequencing library according to the non-Amplified non-Tagging Illumina Cap Analysis of Gene Expression (nAnT-iCAGE) (*23*) by using CAGE library preparation kit (DNAFORM), and sequenced by NextSeq500 platform (Illumina). After discarding sequences with ambiguous (N) and low-quality (Phred<30), the remaining reads were aligned with the human reference genome (GRCh37) by STAR v2.6.0a (*54*) with a guide of known junctions of RefSeq transcripts (*55*). The alignments with mapping quality more than 20 were selected, and their 5’-ends were counted based on the robust set of CAGE peaks identified in a previous study (*56*) and provided at the FANTOM5 web resource (*57*). The read counts were normalized as counts per million (CPM) with relative log expression (RLE) method (*58*). The sequencing results were visualized using Integrative Genomics Viewer (IGV) (*59*). This normalization and subsequent differential analysis were conducted with edgeR (*60*), where FDR<1e-4 were used as a criteria for statistical significance. *p* value of GO term enrichment was assessed with DAVID v6.8 (*61*) using all human genes as a background set, and only the terms that have less than 100 corresponding genes were chosen to exclude very general GO terms such as “biological process”. The obtained CAGE reads are available in the DDBJ Sequence Read Archive (DRA) with accession number (DRA007587).

### Clinical pancreatic tissue samples

The pancreatic tissues used in this study were obtained from patients who underwent surgical treatment at Tokyo Metropolitan Geriatric Hospital (n=47; female, n=25; male, n=22; 62-91 years old; mean age, 74.8±7.0 years old). The present study was conducted in accordance with the principles embodied in the Declaration of Helsinki, 2013, and all experiments were approved by the ethics committees of Tokyo Metropolitan Geriatric Hospital and Institute of Gerontology (permit-#260219).

Tissues were fixed in 10% buffered formalin and then subjected to standard tissue processing and paraffin embedding. The tissues were sliced serially into sections 3 µm thick for hematoxylin and eosin (H&E) and immunohistochemical staining. Pathological specimens were diagnosed by our pathologists based on the World Health Organization Classification of Tumours of the Digestive System (*62*). PanIN lesions were classified as PanIN-1, −2 or −3.

### Immunohistochemistry (IHC) of pancreatic tissue samples

Paraffin-embedded sections (3 µm) were subjected to immunostaining. After deparaffinization, the tissue sections were preheated in HEAT PROCESSOR Solution pH 6 (Nichirei) for 20 min at 100°C. Then, sections were incubated for 5 min at room temperature with Protein Block Serum-Free (Dako). The tissue sections were then incubated with the anti-249T-P pAbs or TpMab-1 mAb (1: 250 in dilution) for 1 hour at room temperature. Endogenous peroxidase activity was blocked by 3% H2O2 for 5 min at room temperature. Sections were incubated with Second antibody (REAL EnVision, Dako). Bound antibodies were detected using diaminobenzidine tetrahydrochloride as the substrate. The sections were then counterstained with Mayer’s hematoxylin. Negative control tissue sections were prepared by omitting the primary antibody. As for the evaluation of immunostaining, proportion of positively stained nucleus of cancer cells were analyzed at magnification x200. For statistical analysis, we divided patients into 2 groups; 249T-P positive and negative groups at 10% cutoff value.

### Statistical analysis of pancreatic tissue sample

The level of significance was set at *p*<0.05 for all analyses. Statistical analyses were performed using the StatView J version 5.0 software package (SAS Institute).

### Clinical HCC samples

A total of 100 HCC patients who received surgery at Kanazawa University Hospital from 2008 to 2013 were enrolled in the study. This study was approved by the Institutional Review Board at Kanazawa University (IRB # 1065) and all patients provided written informed consent.

### IHC of HCC samples

IHC of HCC samples was performed using DAKO Envision+ kits (Agilent Technologies) according to the manufacturer’s instruction. Briefly, formalin-fixed paraffin-embedded tissue slides were deparaffinized, rehydrated, and immediately proceeded for antigen retrieval (120°C for 5 min) using autoclaves and target retrieval solution, citrate pH 6 (Agilent Technologies). Slides were immersed with blocking solution (Agilent Technologies) for 15 min and subsequently replaced with anti-249T-P pAbs diluted at 1:250 with antibody diluent solution (Agilent Technologies). Slides were incubated at 4°C overnight and then washed and visualized with DAB+ substrate chromogen (Agilent Technologies). The patient groups with 249T-P positive and negative were divided at 10% cutoff value.

### Statistical analysis of HCC samples

Kaplan-Meier survival analysis was performed in GraphPad Prism software 6.0 (GraphPad Software). The association of phosphorylation of hTERT threonine 249 and clinicopathologic characteristics was examined with either Student’s tests or *χ*2 tests.

## Acknowledgments

We would like to thank Yasuhiro Murakawa for useful discussions. We are grateful to the Support Unit for Bio-Material Analysis, RIKEN CBS RRC, for technical help with the mass spectrometry analysis.

## Author contributions

M.Y. and Y.A. performed biochemical and cellular experiments. T.Y., Y.M. and S.K. performed immunohistological and statistical analysis of clinical samples. S.S. and M.S. designed and performed *in vitro* kinase assay. M.S.M., H.K. and Y.H. performed CAGE sequencing and the bioinformatics analyses. K.S. and T.K. carried out mass spectrometric protein profiling and the bioinformatics analyses. T.A. and Y.F. designed the experiments for CRISPR-Cas9 gene editing of hTERT. S.Y., M.K.K. and Y.K. generated a phospho-specific monoclonal antibody against hTERT. M.Y., Y.A., T.Y., and K.M. designed the experiments and discussed the interpretation of the results. M.Y., Y.A. and K.M. wrote the manuscript.

## Data and materials availability

All data needed to evaluate the conclusions in the paper are present in the paper and/or the Supplementary Materials. Additional data related to this paper may be requested from the authors. The CAGE reads are available in the DDBJ Sequence Read Archive (DRA) under accession number DRA007587.

## Supplementary Materials

**Fig. S1.** Confirmation of anti-hTERT antibodies used in the study.

**Fig. S2.** Mitotic specific accumulation of hTERT mRNA in HeLa cells.

**Fig. S3.** Confirmation of phosphorylation site of hTERT_191-306 protein.

**Fig. S4.** Validation of the specificity of the anti-249T-P antibodies for IHC staining.

**Fig. S5.** Confirmation of the specificity of anti-239T-P and TpMab-1 antibodies for IHC staining.

**Fig. S6.** Effects of inhibition of CDK1 activity on *FOXO4* expression.

**Fig. S7.** A model for regulation of *FOXO4* expression via phosphorylation of hTERT at T249.

**Table S1.** MS data of hTERT from in vitro kinase assay.

**Table S2.** Clinicopathological analysis of the hTERT T249 phosphorylation in pancreatic cancer.

**Table S3.** Clinicopathological analysis of the hTERT T249 phosphorylation in liver cancer.

**Table S4.** GO analysis of genes differentially expressed in T249A-CRISPR cells (FDR < 0.01).

**Table S5.** Sequences of primers, siRNAs and oligos used in the work.

**Supplementary Data S1.** MS data of hTERT isolated from 293T or HeLa cells synchronized to mitotic phase.

