## Supplemental data for "CDK1 dependent phosphorylation of hTERT contributes to cancer progression"

**Supplementary Materials**

**Fig. S1.** Confirmation of anti-hTERT antibodies used in the study.

**Fig. S2.** Mitotic specific accumulation of hTERT mRNA in HeLa cells.

**Fig. S3.** Confirmation of phosphorylation site of hTERT_191-306 protein.

**Fig. S4.** Validation of the specificity of the anti-249T-P antibodies for IHC staining.

**Fig. S5.** Confirmation of the specificity of anti-239T-P and TpMab-1 antibodies for IHC staining.

**Fig. S6.** Effects of inhibition of CDK1 activity on *FOXO4* expression.

**Fig. S7.** A model for regulation of *FOXO4* expression via phosphorylation of hTERT at T249.

**Table S1.** MS data of hTERT from in vitro kinase assay.

**Table S2.** Clinicopathological analysis of the hTERT T249 phosphorylation in pancreatic cancer.

**Table S3.** Clinicopathological analysis of the hTERT T249 phosphorylation in liver cancer.

**Table S4.** GO analysis of genes differentially expressed in T249A-CRISPR cells (FDR < 0.01).

**Table S5.** Sequences of primers, siRNAs and oligos used in the work.

**Supplementary Data S1.** MS data of hTERT isolated from 293T or HeLa cells synchronized to mitotic phase.

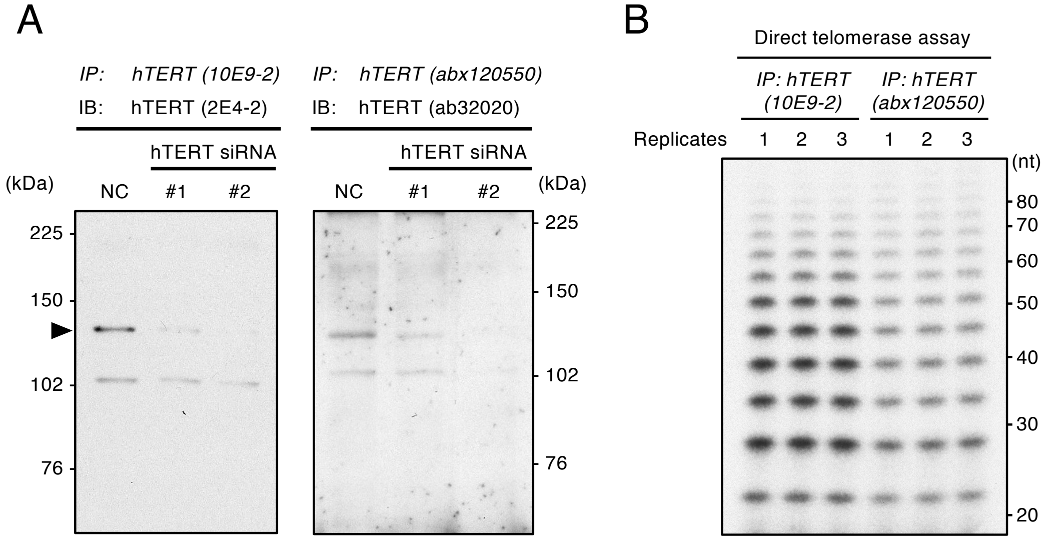


**Figure S1** Confirmation of anti-hTERT antibodies used in the study
**(A)** The endogenous hTERT proteins were immunoprecipitated with anti-hTERT mouse mAb (clone 10E9-2) or anti-hTERT sheep pAbs (abx120550) from HeLa cells transfected with two different siRNAs specific for hTERT or siNC followed by nocodazole treatment. The proteins were detected by anti-hTERT mouse mAb (clone 2E4-2) or anti-hTERT rabbit mAb (ab32020). **(B)** Direct telomerase assay using hTERT proteins immunoprecipitated with anti-hTERT mouse mAb (clone 10E9-2) or anti-hTERT sheep pAbs (abx120550) from HeLa cells.

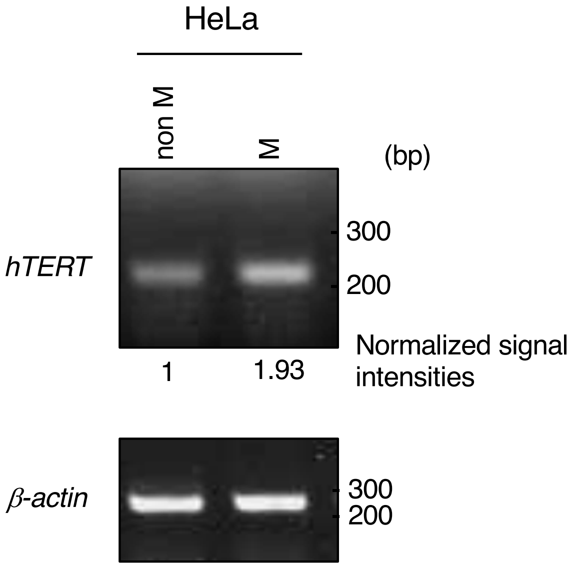


**Figure S2** Mitotic specific accumulation of hTERT mRNA in HeLa cells
RT-PCR was performed to confirm the expression of *hTERT* mRNA in HeLa cells treated with DMSO (non M) or nocodazole (M). The normalized signal intensities with *β-actin* are noted below the panel.

hTERT_191-306

191 206
SGPRRRLGCE RAWNH**S**VREA GVPLGLPAPG

249
ARRRGGSASR SLPLPKRPRR GAAPEPER**T**P

274
VGQGSWAHPG RTRGPSDRGF CVV**S**PARPAE

283 306
EA**T**SLEGALS GTRHSHPSVG RQHHAG

**Figure S3** Confirmation of phosphorylation site of hTERT_191-306 protein

The recombinant hTERT fragment proteins (hTERT_191-306) were phosphorylated by CDK1-cyclinB or IKK2_2-664 *in vitro* and analyzed by MS to confirm the phosphorylation sites. Threonine residue denoted with red letter was specifically phosphorylated by CDK1-cyclinB. Blue letter indicates threonine residue phosphorylated by IKK2_2-664 and green letters indicate serine residues phosphorylated by both CDK1-cyclinB and IKK2_2-664. See the details in **Table S1**.

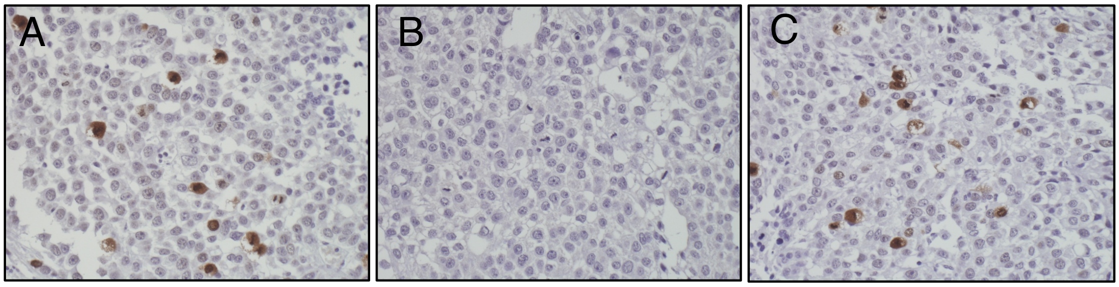


**Figure S4** Validation of the specificity of the anti-249T-P antibodies for IHC staining

**(A)** Paraffin embedded sections of Huh7 xenografts were stained with anti-249T-P antibodies. **(B** and **C)** Paraffin embedded sections of Huh7 xenografts were incubated with 0.4 $\mu$g/mL of phosphopeptides (**B**) or nonphosphopeptides (**C**) simultaneously with the same concentration of anti-249T-P pAbs.

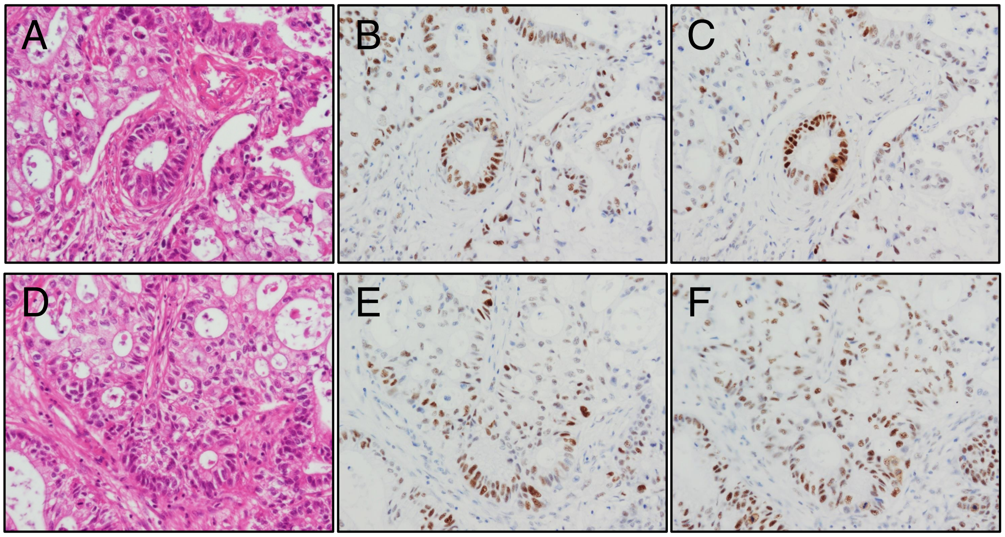


**Figure S5** Confirmation of the specificity of anti-239T-P and TpMab-1 antibodies for IHC staining

**(A-F)** Hematoxilyn and elosin (HE) staining (**A** and **D**), IHC staining with anti-249T-P pAbs (**B** and **E**) and with TpMab-1 antibody (**C** and **F**) in serial sections (**A-C** and **D-F**) prepared from the identical pancreatic cancer lesions.

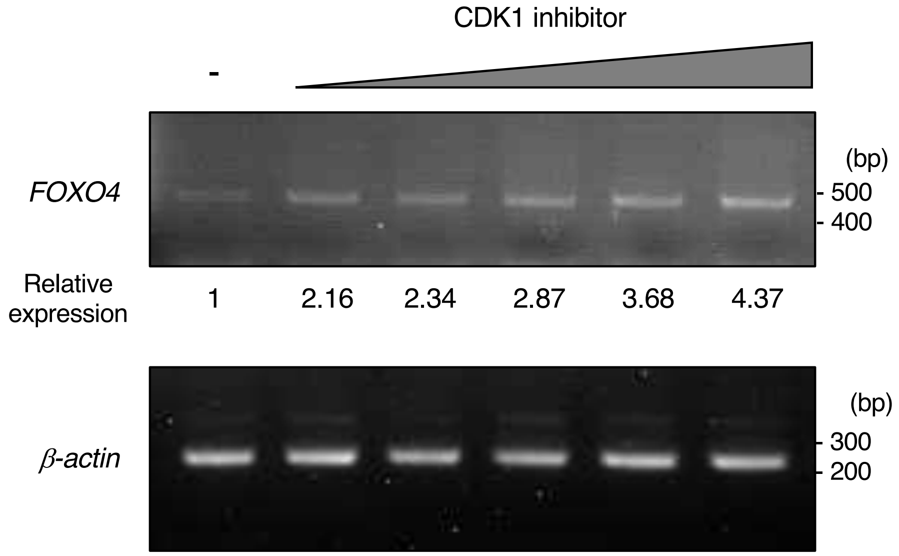


**Figure S6** Effects of inhibition of CDK1 activity on *FOXO4* expression
Increase of *FOXO4* mRNAs were detected by RT-PCR (upper panel). Total RNAs were extracted from HeLa cells treated with CDK1 inhibitor, RO-3306 (0, 0.15625, 0.3125, 0.625, 1.25, 2.5 $\mu$M) and nocodazole (100 ng/mL). *β-actin* was used as an internal control (lower panel).

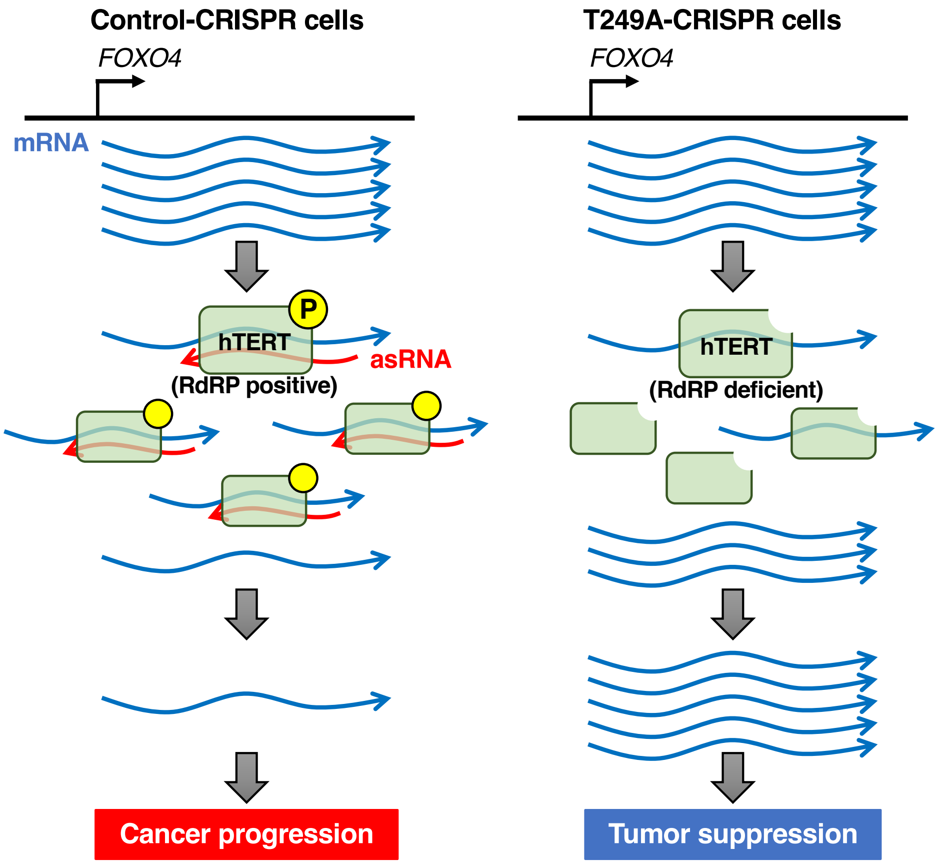


**Figure S7** A model for regulation of *FOXO4* expression via phosphorylation of hTERT at T249

In Control-CRISPR cells, hTERT proteins have phosphorylation-dependent RdRP activity and synthesize asRNAs from *FOXO4* mRNAs. These RNAs form double-stranded RNAs and might be degraded. In T249A-CRISPR cells, RdRP-deficient hTERT-T249A proteins interact with less *FOXO4* mRNAs. Without producing asRNA, more *FOXO4* mRNAs are retained in the T249A-CRISPR cells and increase of *FOXO4* expression cause tumor suppression.

**Table S1**, MS data of hTERT from in vitro kinase assay

hTERT.191-306_Phosphorylation by CDK1-CycB

| Description | Score | Coverage | # Proteins | # Unique Peptides | # Peptides | # PSMs | # AAs | MW [kDa] | calc. pI |
|---|---|---|---|---|---|---|---|---|---|
| Telomerase reverse transcriptase OS=Homo sapiens GN=TERT PE=1 SV=1 - [TERT_HUMAN] | 781.66 | 7.77 | 1 | 12 | 12 | 46 | 1132 | 126.9 | 10.52 |

| A2 | Sequence | # PSMs | # Proteins | # Protein Groups | Protein Group Accessions | 1 | ΔCn | phosphoRS Site Probabilities | q-Value | PEP | IonScore | Exp Value | Charge | MH+ [Da] | ΔM [ppm] | RT [min] | # Missed Cleavages |
|---|---|---|---|---|---|---|---|---|---|---|---|---|---|---|---|---|---|
| High | TPVGQGSWAHPGR | 2 | 1 | 1 | O14746 | | 0.0000 | | 0 | 0.0001284 | 68 | 1.11476E-06 | 2 | 1349.67143 | 0.34 | 13.45 | 0 |
| High (hTERT_274S) | GFcVVsPARPAEEATSLE | 3 | 1 | 1 | O14746 | C3(Carbamidomethyl); S | 0.0000 | S(6): 100.0; T(15): 0.0; S( | 0 | 0.00005126 | 61 | 7.83771E-06 | 3 | 2642.23880 | 4.03 | 31.58 | 0 |
| High | EAGVPLGLPAPGAR | 12 | 1 | 1 | O14746 | | 0.0000 | | 0 | 0.0002435 | 59 | 3.52226E-06 | 2 | 1304.73235 | 0.11 | 25.07 | 0 |
| High | GFcVVSPARPAEEATSLE | 1 | 1 | 1 | O14746 | C3(Carbamidomethyl) | 0.0000 | | 0 | 0.0002302 | 48 | 0.000118526 | 3 | 2562.26200 | 0.07 | 28.50 | 0 |
| High | GPSDRGFcVVSPARPAE | 2 | 1 | 1 | O14746 | C8(Carbamidomethyl) | 0.0000 | | 0 | 0.01813 | 44 | 0.000280259 | 4 | 3074.51333 | 5.58 | 27.51 | 1 |
| High | RGAAPEPER | 1 | 1 | 1 | O14746 | | 0.0000 | | 0 | 0.02462 | 43 | 0.000298225 | 2 | 982.50737 | 0.86 | 7.16 | 1 |
| **High (hTERT_249T)** | **GAAPEPERtPVGQGSW** | **1** | **1** | **1** | **O14746** | **T9(Phospho)** | **0.0000** | **T(9): 100.0;** S(15): 0.0 | **0** | **0.007542** | **42** | **0.000698864** | **4** | **2237.02480** | **-0.03** | **15.88** | **1** |
| High (hTERT_274S) | GPSDRGFcVVsPARPAE | 8 | 1 | 1 | O14746 | C8(Carbamidomethyl); S | 0.0000 | S(3): 0.0; S(11): 100.0; T( | 0 | 0.0002018 | 42 | 0.000769782 | 4 | 3154.46694 | 1.41 | 29.31 | 1 |
| **High (hTERT_249T)** | **tPVGQGSWAHPGR** | **1** | **1** | **1** | **O14746** | **T1(Phospho)T1(Phosp** | **0.0000** | **T(1): 100.0;** S(7): 0.0 | **0** | **0.09146** | **36** | **0.001734836** | **2** | **1429.63677** | **-0.37** | **15.27** | **0** |
| High | AWNHSVREAGVPLGLP | 4 | 1 | 1 | O14746 | | 0.0000 | | 0 | 0.00304 | 29 | 0.003476626 | 3 | 2155.15390 | 0.86 | 22.50 | 1 |
| High | SLPLPKRPR | 3 | 1 | 1 | O14746 | | 0.0000 | | 0 | 0.02031 | 23 | 0.004720304 | 3 | 1063.67411 | 0.54 | 12.70 | 1 |
| High | EAGVPLGLPAPGARR | 2 | 1 | 1 | O14746 | | 0.0000 | | 0 | 0.00247 | 21 | 0.01783279 | 3 | 1460.83353 | 0.14 | 21.23 | 1 |
| High (hTERT_206S) | AWNHsVREAGVPLGLP | 1 | 1 | 1 | O14746 | S5(Phospho) | 0.0000 | S(5): 100.0 | 0 | 0.1816 | 20 | 0.081900648 | 3 | 2235.11753 | -0.38 | 24.55 | 1 |
| High | GAAPEPERTPVGQGSW | 2 | 1 | 1 | O14746 | | 0.0000 | | 0 | 0.0787 | 15 | 0.258115362 | 5 | 2157.05717 | -0.64 | 15.74 | 1 |
| High | LGcERAWNHSVR | 1 | 1 | 1 | O14746 | C3(Carbamidomethyl) | 0.0000 | | 0 | 0.06023 | 10 | 0.749015362 | 3 | 1484.71708 | -0.36 | 11.67 | 1 |
| Medium | GAAPEPER | 1 | 1 | 1 | O14746 | | 0.0000 | | 0.022 | 0.2064 | 35 | 0.002318774 | 2 | 826.40520 | -0.26 | 7.43 | 0 |
| Medium | AWNHSVR | 1 | 1 | 1 | O14746 | | 0.0000 | | 0.028 | 0.2521 | 26 | 0.007735331 | 2 | 869.43804 | 0.40 | 7.98 | 0 |

hTERT.191-306_Phosphorylation by IKK2

| Description | Score | Coverage | # Proteins | # Unique Peptides | # Peptides | # PSMs | # AAs | MW [kDa] | calc. pI |
|---|---|---|---|---|---|---|---|---|---|
| Telomerase reverse transcriptase OS=Homo sapiens GN=TERT PE=1 SV=1 - [TERT_HUMAN] | 1019.70 | 7.33 | 1 | 9 | 9 | 75 | 1132 | 126.9 | 10.52 |

| A2 | Sequence | # PSMs | # Proteins | # Protein Groups | Protein Group Accessions | Modifications | ΔCn | phosphoRS Site Probabilities | q-Value | PEP | IonScore | Exp Value | Charge | MH+ [Da] | ΔM [ppm] | RT [min] | # Missed Cleavages |
|---|---|---|---|---|---|---|---|---|---|---|---|---|---|---|---|---|---|
| High | TPVGQGSWAHPGR | 5 | 1 | 1 | O14746 | | 0.0000 | | 0 | 1.392E-14 | 56 | 2.11052E-05 | 3 | 1349.66883 | -1.59 | 13.96 | 0 |
| High | GFcVVSPARPAEEATSLE | 8 | 1 | 1 | O14746 | C3(Carbamidomethyl) | 0.0000 | | 0 | 6.228E-16 | 49 | 8.53574E-05 | 3 | 2562.27518 | 5.22 | 28.64 | 0 |
| High (hTERT_206S) | AWNHsVREAGVPLGLP | 42 | 1 | 1 | O14746 | S5(Phospho) | 0.0000 | S(5): 100.0 | 0 | 4.6E-15 | 47 | 0.000146669 | 3 | 2235.11972 | 0.60 | 25.11 | 1 |
| High | GPSDRGFcVVSPARPAE | 2 | 1 | 1 | O14746 | C8(Carbamidomethyl) | 0.0000 | | 0 | 3.66009E-16 | 47 | 0.000192597 | 4 | 3074.50063 | 1.46 | 27.01 | 1 |
| High | RGAAPEPER | 1 | 1 | 1 | O14746 | | 0.0000 | | 0 | 1.862E-14 | 43 | 0.000298225 | 2 | 982.50682 | 0.30 | 7.16 | 1 |
| High (hTERT_283T) | GPSDRGFcVVSPARPAE | 2 | 1 | 1 | O14746 | C8(Carbamidomethyl); T | 0.0000 | S(3): 0.0; S(11): 0.0; T(20 | 0 | 3.976E-11 | 39 | 0.001401446 | 4 | 3154.46181 | -0.21 | 27.83 | 1 |
| High | EAGVPLGLPAPGAR | 5 | 1 | 1 | O14746 | | 0.0000 | | 0 | 0.000004751 | 37 | 0.000625746 | 2 | 1304.73332 | 0.86 | 24.45 | 0 |
| High | GAAPEPERTPVGQGSW | 5 | 1 | 1 | O14746 | | 0.0000 | | 0 | 1.655E-07 | 32 | 0.005081198 | 4 | 2157.05922 | 0.32 | 15.96 | 1 |
| High | EAGVPLGLPAPGARR | 1 | 1 | 1 | O14746 | | 0.0000 | | 0 | 1.789E-11 | 26 | 0.004211957 | 3 | 1460.83820 | 3.34 | 21.00 | 1 |
| High | SLPLPKRPR | 3 | 1 | 1 | O14746 | | 0.0000 | | 0 | 0.008781 | 26 | 0.002666674 | 3 | 1063.67411 | 0.54 | 12.72 | 1 |
| High (hTERT_274S) | GFcVVsPARPAEEATSLE | 1 | 1 | 1 | O14746 | C3(Carbamidomethyl); S | 0.0000 | S(6): 96.6; T(15): 1.1; S(1 | 0 | 0.003605 | 12 | 0.599837122 | 3 | 2642.22965 | 0.57 | 28.26 | 0 |

**Table S2**, Clinicopathological analysis of the hTERT T249 phosphorylation in pancreatic cancer

| | | 249T-P | | |
|---|---|---|---|---|
| | | Positive (n=26) | Negative (n=21) | *p* value |
| Age (yr, mean ± SE) | | 74.8 ± 1.3 | 74.8 ± 1.7 | 0.9972 |
| Gender | Female | 17 | 8 | 0.0623 |
| | Male | 9 | 13 | |
| Histological differentiation | Well | 10 | 12 | 0.3221 |
| | Moderate | 13 | 7 | |
| | Poor | 0 | 1 | |
| | Adenoaquamous carcinoma | 3 | 1 | |
| Primary tumor (T) | T0 | 1 | 0 | 0.4588 |
| | T1 | 2 | 0 | |
| | T2 | 1 | 1 | |
| | T3 | 22 | 20 | |
| Lymph nodes (N) | N0 | 6 | 12 | 0.0169* |
| | N1 | 20 | 9 | |
| Metastasis (M) | M0 | 25 | 20 | 0.8771 |
| | M1 | 1 | 1 | |

Asterisk indicates statistically significant values ($p<0.05$).

**Table S3**, Clinicopathological analysis of the hTERT T249 phosphorylation in liver cancer

| | | 249T-P | | |
|---|---|---|---|---|
| | | Positive (n = 29) | Negative (n = 71) | *p* value |
| Age (yr, mean ± SE) | | 65.7 ± 2.2 | 64.8 ± 1.2 | 0.7155 |
| Gender | Female | 4 | 24 | 0.0432* |
| | Male | 25 | 47 | |
| Etiology | HBV | 5 | 16 | 0.7889 |
| | HCV | 11 | 28 | |
| | B+C | 0 | 1 | |
| | other | 13 | 26 | |
| AFP (ng/ml, mean ± SE) | | 60.5 ± 24.1 | 2,311 ± 1,033 | 0.165 |
| Histological grade | Well | 2 | 22 | 0.0122* |
| | Moderate | 21 | 44 | |
| | Poor | 6 | 5 | |
| Tumor size | <5cm | 22 | 52 | 0.7862 |
| | >5cm | 7 | 19 | |
| BCLC stage | A | 15 | 46 | 0.3183 |
| | B | 7 | 16 | |
| | C | 7 | 9 | |

Asterisk indicates statistically significant values ($p<0.05$).
Abbreviation: AFP, alphafetoprotein. BCLC, Barcelona Clinic Liver Cancer

**Table S4**, GO analysis of genes differentially expressed in T249A-CRISPR cells (FDR < 0.01).

| UP/DOWN | Term | # of genes | P value | Fold Enrichment | FDR |
|---|---|---|---|---|---|
| UP | GO:0042254~ribosome biogenesis | 82 | 1.87E-21 | 3.213667394 | 3.70E-18 |
| | GO:0006364~rRNA processing | 66 | 1.86E-17 | 3.221243362 | 3.67E-14 |
| | GO:0016072~rRNA metabolic process | 66 | 7.31E-17 | 3.139545161 | 2.22E-13 |
| | GO:0034470~ncRNA processing | 80 | 1.07E-14 | 2.555524484 | 2.11E-11 |
| | GO:0006281~DNA repair | 89 | 2.03E-12 | 2.204682314 | 4.02E-09 |
| | GO:0008380~RNA splicing | 73 | 4.61E-12 | 2.396043784 | 9.11E-09 |
| | GO:0006412~translation | 98 | 3.74E-11 | 2.007242885 | 7.38E-08 |
| | GO:0006397~mRNA processing | 78 | 1.09E-10 | 2.174230465 | 2.16E-07 |
| | GO:0022618~ribonucleoprotein complex assembly | 45 | 7.41E-10 | 2.786817531 | 1.46E-06 |
| | **GO:0044770~cell cycle phase transition** | 84 | 8.92E-10 | 2.016154645 | 1.76E-06 |
| | GO:0016458~gene silencing | 50 | 9.89E-10 | 2.594665422 | 1.96E-06 |
| | GO:0071826~ribonucleoprotein complex subunit organization | 46 | 1.23E-09 | 2.708225667 | 2.43E-06 |
| | **GO:0044772~mitotic cell cycle phase transition** | 79 | 2.89E-09 | 2.017882404 | 5.71E-06 |
| | GO:0000375~RNA splicing, via transesterification reactions | 55 | 3.24E-09 | 2.383153092 | 6.40E-06 |
| DOWN | GO:0000184~nuclear-transcribed mRNA catabolic process, nonsense | 36 | 1.65E-17 | 5.788847712 | 3.23E-14 |
| | GO:0006614~SRP-dependent cotranslational protein targeting to mem | 32 | 2.11E-17 | 6.592854338 | 4.13E-14 |
| | GO:0006613~cotranslational protein targeting to membrane | 33 | 2.32E-17 | 6.33682116 | 4.53E-14 |
| | GO:0006413~translational initiation | 43 | 1.87E-16 | 4.499884707 | 4.33E-13 |
| | GO:0045047~protein targeting to ER | 32 | 2.71E-16 | 6.085711697 | 4.33E-13 |
| | GO:0070972~protein localization to endoplasmic reticulum | 35 | 5.86E-16 | 5.408200824 | 1.09E-12 |
| | GO:0072599~establishment of protein localization to endoplasmic retic | 32 | 9.89E-16 | 5.860314967 | 1.95E-12 |
| | GO:0006612~protein targeting to membrane | 39 | 2.17E-13 | 4.059810303 | 4.25E-10 |
| | GO:0019080~viral gene expression | 37 | 6.88E-12 | 3.811493914 | 1.35E-08 |
| | GO:0019058~viral life cycle | 61 | 7.93E-12 | 2.651631525 | 1.55E-08 |
| | GO:0000956~nuclear-transcribed mRNA catabolic process | 38 | 1.33E-11 | 3.648472789 | 2.60E-08 |
| | GO:0006402~mRNA catabolic process | 39 | 2.74E-11 | 3.49033465 | 5.36E-08 |
| | GO:0044033~multi-organism metabolic process | 38 | 4.24E-11 | 3.512081283 | 8.29E-08 |
| | GO:0006401~RNA catabolic process | 41 | 7.21E-11 | 3.256711179 | 1.41E-07 |
| | GO:0019083~viral transcription | 34 | 1.13E-10 | 3.71531018 | 2.21E-07 |
| | GO:0006518~peptide metabolic process | 86 | 1.45E-10 | 2.079408826 | 2.85E-07 |
| | GO:0090150~establishment of protein localization to membrane | 51 | 1.60E-10 | 2.741050853 | 3.13E-07 |
| | GO:0043043~peptide biosynthetic process | 74 | 3.21E-10 | 2.194323333 | 6.28E-07 |
| | GO:0043604~amide biosynthetic process | 79 | 3.54E-10 | 2.122970758 | 6.92E-07 |
| | GO:0006412~translation | 70 | 2.03E-09 | 2.159905477 | 3.97E-06 |

**Table S5**, Sequences of primers, siRNAs and oligos used in the work

Mutagenic primers

| | |
|---|---|
| T249A | AGCCGGAGCGGGCGCCCGTTGGG |
| | CCCAACGGGCGCCCGCTCCGGCT |
| T249E | TGAGCCGGAGCGGGAGCCCGTTGGGCAG |
| | CTGCCCAACGGGCTCCCGCTCCGGCTCA |

PCR primers for qPCR and RT-PCR

| | |
|---|---|
| hTERT_F | CGGAAGAGTGTCTGGAGCAA |
| hTERT_R | GGATGAAGCGGAGTCTGGA |
| hLINE1_F | TTGGAAAACACTCTGCAGGATATTAT |
| hLINE1_R | TTGGCCTGCCTTGCTAGATT |
| hGAPDH_F | GAAGGTGAAGGTCGGAGTCA |
| hGAPDH_R | GAAGATGGTGATGGGATTTC |
| Alpha-satellite_RT | CCGTAAAACGACGGCCAGCTTCTGTCTAGTTTTTATGTGAAGATA |
| Alpha-satellite_F | CATTCTCAGAAACTTCTTTGTGATGTG |
| Alpha-satellite_R | CCGTAAAACGACGGCCAG |
| RMRP-S_RT | AGCCGCGCTGAGAATGAG |
| RMRP-AS_RT | GTGCTGAAGGCCTGTATCCT |
| RMRP_F | TGCTGAAGGCCTGTATCCT |
| RMRP_R | TGAGAATGAGCCCCGTGT |
| β-actin_F | CAAGAGATGGCCACGGCTGCT |
| β-actin_R | TCCTTCTGCATCCTGTCGGCA |
| FOXO4_RT | AAGTGTCAGTCGCTTCTCCG |
| FOXO4_F | AAAAAGTGCTTCGCTCGGC |
| FOXO4_R | GCTGGTTAGCGATCTCTGGT |
| FOXO4 asRNA_RT | GGGATACAGTGCCTCAGGTTT |
| FOXO4 asRNA_F | AAGTGTCAGTCGCTTCTCCG |
| FOXO4 asRNA_R | AAAAAGTGTCTTCGCTCGGC |
| RNaseP_F | GTCACTCCACTCCCATGTCC |
| RNaseP_R | AATTGGGTTATGAGGTCCCC |

siRNAs

| | |
|---|---|
| TERT siRNA #1 | GUGUCUGUGCCCGGGAGAATT |
| | UUCUCCCGGGCACAGACACTT |
| TERT siRNA #2 | GCAUUGGAAUCAGACAGCATT |
| | UGCUGUCUGAUUCCAAUGCTT |

Oligos for CRISPR

| | |
|---|---|
| guide RNA sequence | AGCCGGAGCGGacgCCCGTT GGG |
| Insert sequence of donor plasmid (pBluescript/SalI-NotI) for T249A | gtcgacGTCCCCCTGGGCCTGCCAGCCCCGGGTGCGAGGAGGCGCGGGGGCAGTGCCAGCCGA AGTCTGCCGTTGCCCAAGAGGCCCAGGCGTGGCGCTGCCCCTGAGCCGGAGCGGgccCCCGTT GGGCAGGGGTCCTGGGCCCACCCGGGCAGGACGCGTGGACCGAGTGACCGTGGTTTCTGTGT GGTGTCACCTGCCAGACCCGCCGAAGAAGCCACCTCTTTGGcggccgc |
| Donor oligo for T249E | GCCGAAGTCTGCCGTTGCCCAAGAGGCCCAGGCGTGGCGCTGCCCCTGAGCCGGAGCGGgaa CCgGTTGGGCAGGGGTCCTGGGCCCACCCGGGCAGGACGCGTGGACCGAGTGACCGTG |
| CRISPR PCR_F | AGCTACCTGCCCAACACGGT |
| CRISPR PCR_R for T249A | CGCACGCTCATCTTCCACGT |
| CRISPR PCR_R for T249E | CACACAGAAACCACGGTCAC |

FOXO4 shRNAs

| | |
|---|---|
| shFOXO4-1 (TRCN0000010291) | CCGGCACTTAGGCTTTGTAGCAAGACTCGAGTCTTGCTACAAAGCCTAAGTGTTTTTG |
| shFOXO4-2 (TRCN0000039720) | CCGGCCAGCTTCAGTCAGCAGTTATCTCGAGATAACTGCTGACTGAAGCTGGTTTTTG |
